# Nemertean, brachiopod and phoronid neuropeptidomics reveals ancestral spiralian signalling systems

**DOI:** 10.1101/2021.03.03.433790

**Authors:** Daniel Thiel, Luis A. Yañez Guerra, Mirita Franz-Wachtel, Andreas Hejnol, Gáspár Jékely

**Affiliations:** Living Systems Institute, University of Exeter, Stocker Road, Exeter, UK; Eberhard Karls Universität Tübingen, Interfaculty Institute for Cell Biology, Tübingen, Germany; University of Bergen, Department of Biological Sciences, 5006 Bergen, Norway

**Author notes:** Equal contribution (Daniel Thiel, Luis A. Yañez Guerra).

**Keywords:** RFamide, Agatoxin-like peptide, Pleurin, APGWamide, Neuropeptide, GPCRs, Trochozoa, GnRH, Vasopressin, GPR139

## Abstract

Neuropeptides are diverse signalling molecules in animals commonly acting through G-protein coupled receptors (GPCRs). Neuropeptides and their receptors underwent extensive diversification in bilaterians and the relationships of many peptide-receptor systems have been clarified. However, we lack a detailed picture of neuropeptide evolution in lophotrochozoans as in-depth studies only exist for molluscs and annelids. Here we analyse peptidergic systems in Nemertea, Brachiopoda and Phoronida. We screened transcriptomes from thirteen nemertean, six brachiopod and four phoronid species for proneuropeptides and neuropeptide GPCRs. With mass spectrometry from the nemertean *Lineus longissimus*, we validated several predicted peptides and identified novel ones. Molecular phylogeny combined with peptide-sequence and gene-structure comparisons allowed us to comprehensively map spiralian neuropeptide evolution. We found most mollusc and annelid peptidergic systems also in nemerteans, brachiopods and phoronids. We uncovered previously hidden relationships including the orthologies of spiralian CCWamides to arthropod agatoxin-like peptides and of mollusc APGWamides to RGWamides from annelids, with orthologues systems in nemerteans, brachiopods and phoronids. We found that pleurin neuropeptides previously only found in molluscs are also present in nemerteans and brachiopods. We also identified cases of gene family duplications and losses. These include a protostome-specific expansion of RFamide/Wamide signalling, a spiralian expansion of GnRH-related peptides, and duplications of vasopressin/oxytocin before the divergence of brachiopods, phoronids and nemerteans. This analysis expands our knowledge of peptidergic signalling in spiralians and protostomes. Our annotated dataset of nearly 1,300 proneuropeptide sequences and 600 GPCRs presents a useful resource for further studies of neuropeptide signalling in protostomes.

## Introduction

Neuropeptides are a diverse group of neuronal signaling molecules found in most animals (Jekely, 2013; Mirabeau and Joly, 2013; Elphick, Mirabeau and Larhammar, 2018; Thiel *et al*., 2018; Quiroga Artigas *et al*., 2020). Most mature neuropeptides consist of 2-40 amino acids and derive from longer proneuropeptide (pNP) precursor sequences. Precursor sequences can be a few hundred amino acids long containing one or multiple neuropeptides. The active peptides in a pNP can be interspersed with other, often non-conserved peptides (intersequences) with unknown biological function (Jekely, 2013; Mirabeau and Joly, 2013; Christie, 2017; Veenstra, 2019; Takahashi, 2020). Neuropeptide sequences in pNPs are generally flanked by basic residues for enzymatic cleavage (Veenstra, 2000; Hook *et al.*, 2008). The phylogenomic analysis of neuropeptides can be challenging due to the short length of the active peptides and often high degrees of divergence and intersequences are usually even less conserved between species. Nevertheless, it has been possible to reconstruct the deep evolutionary history of most bilaterian peptidergic systems by the combined analysis of pNPs and neuropeptide receptors (Jekely, 2013; Mirabeau and Joly, 2013; Tian *et al.*, 2016; Thiel *et al.*, 2018).

Most neuropeptides activate G-protein coupled receptors (GPCRs) that belong either to the class A (rhodopsin type) or class B (secretin type) of GPCRs (Caers *et al.*, 2012; Frooninckx *et* al., 2012; Mirabeau and Joly, 2013; Foster *et al.*, 2019). The longer (several hundred amino acids) and more conserved sequence of GPCRs make phylogenetic analyses feasible when reconstructing the evolution of peptidergic systems (Jekely, 2013; Mirabeau and Joly, 2013). Another useful approach is to compare the exon-intron structure of pNPs (Mair *et al.*, 2000; Yañez-Guerra *et al.*, 2020; Zhang *et al.*, 2020). In several cases, the overall organisation of orthologous pNP genes can be conserved (e.g. a region coding for the signal peptide followed by a single peptide and a C-terminal Cys-containing domain) (Semmens *et al.*, 2015; Liutkeviciute *et al.*, 2016). Furthermore, an increased sampling across taxa has often helped to clarify relationships, revealing hidden orthologs or lineage specific losses, gains and divergences (Jekely, 2013; Mirabeau and Joly, 2013; Semmens *et al.*, 2016; Martín-Durán *et* al., 2017; Thiel *et al.*, 2018; Yañez-Guerra *et al.*, 2020). Some lineages can retain signalling systems that had been lost in related species. For example, corazonin, luqin and short neuropeptide F are widespread neuropeptides in protostomes but not detectable in vertebrates. These pNPs and their receptors are, however, present in ambulacrarians like the hemichordate *Saccoglossus kovalevski* or the starfish *Asterias rubens*, indicating that they represent ancestral bilaterian signaling systems (Tian *et al.*, 2016; Yañez-Guerra *et al.*, 2018, 2020; Yañez-Guerra and Elphick, 2020).

Among the lophotrochozoans, a major and diverse clade of bilaterians, the study of neuropeptides has mostly been limited to molluscs (Veenstra, 2010; Stewart *et al.*, 2014; Adamson *et al.*, 2015; Bose *et al.*, 2017; Oliveira *et al.*, 2019) and annelids (Veenstra, 2011; Conzelmann, Williams, Krug, *et al.*, 2013; Kerbl *et al.*, 2017). Neuropeptide receptors are also known from molluscs and annelids (Conzelmann, Williams, Tunaru, *et al.*, 2013; Jekely, 2013; Mirabeau and Joly, 2013; Bauknecht and Jékely, 2015; Schmidt *et al.*, 2018). These studies have shown that lophotrochozoans possess a relatively conserved neuropeptide complement with evidence for many ancestral bilaterian peptidergic systems (Conzelmann, Williams, Krug, *et al.*, 2013; Oliveira *et al.*, 2019). In addition, molluscs and annelids possess neuropeptide duplications and a few seemingly unique neuropeptides specific to only one of these clades. The neuropeptide complement of other lophotrochozoans – or spiralians in general – has not been sampled thoroughly (Saberi *et al.*, 2016; Oliveira *et al.*, 2019) or was limited to single peptides (Thiel *et al.*, 2017, 2019). A recent study on the evolution of neuropeptides in molluscs surveyed other lophotrochozoans, including nemerteans, brachiopods and phoronids, but only with limited taxonomic sampling (Oliveira *et al.*, 2019). The results failed to recover several ancestral neuropeptides known from molluscs and annelids, raising the question whether this is due to extensive losses during lophotrochozoan evolution.

Here we present a comprehensive bioinformatic survey of the neuropeptide and neuropeptide GPCR complement of nemerteans, brachiopods and phoronids. We investigated several species in each group and were able to identify many previously undetected families. To verify cleavage patterns and the predicted active peptides, we complemented the in silico analysis with a mass spectrometric screen in a nemertean. This enabled us to reconstruct a deeply-sampled complement of peptidergic systems in these organisms and to clarify major evolutionary patterns in lophotrochozoan neuropeptide evolution. We present evidence for protostome-specific expansions of signalling systems, the orthology of some arthropod and spiralian neuropeptides, the lophotrochozoan ancestry of neuropeptides that have been described as clade specific, and describe the evolution of neuropeptide paralogs in different spiralian groups.

## Results

### Bioinformatic identification of peptidergic signalling systems in nemerteans, phoronids and brachiopods

For a comprehensive characterisation of neuropeptide signalling in trochozoans, we searched for neuropeptide precursors and neuropeptide GPCRs in transcriptomes of 13 nemertean, six brachiopod and four phoronid species and analysed them in an evolutionary context (fig. 1).

**Figure 1.**
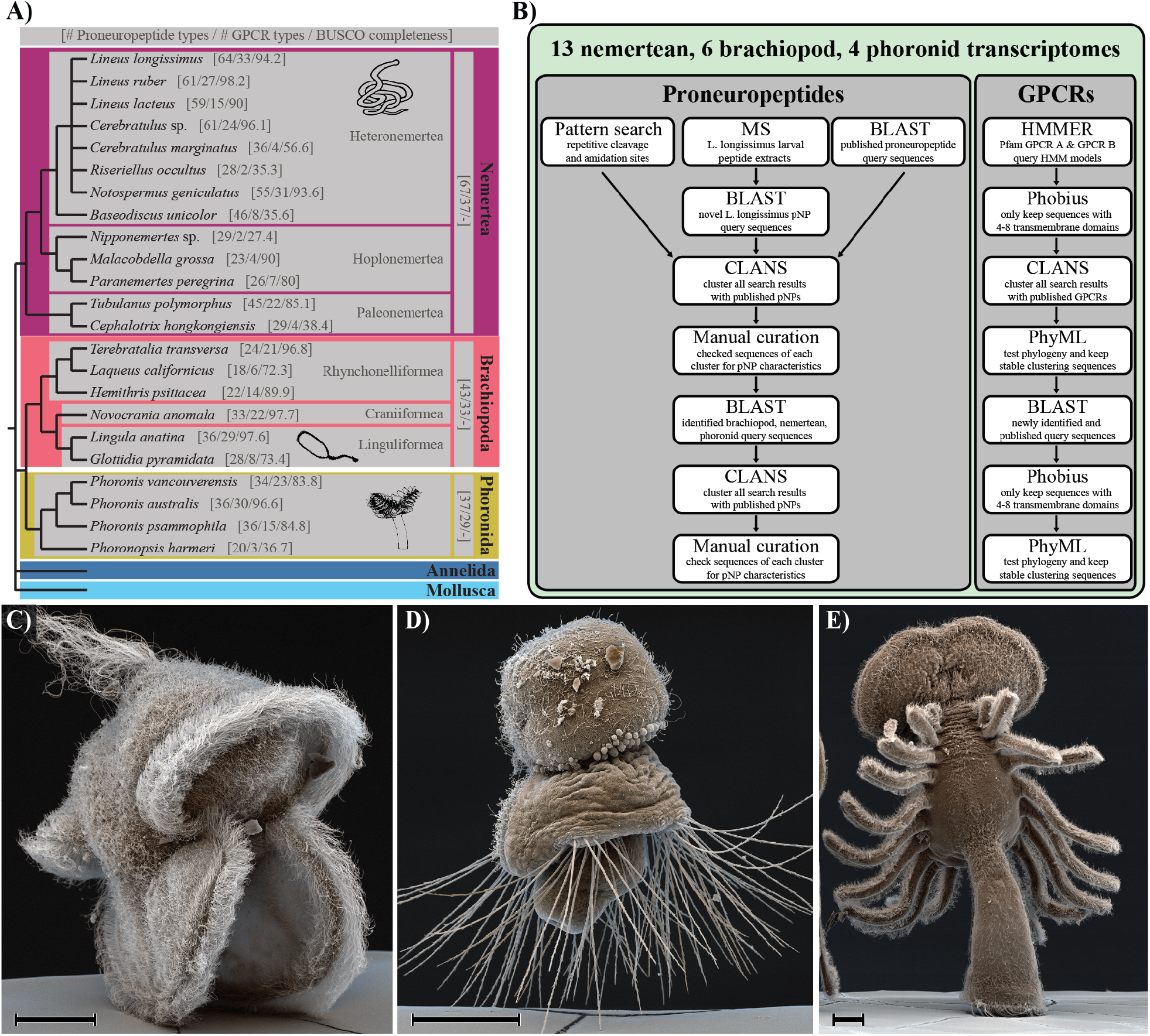
Investigated taxa and pipeline for the identification of peptidergic signaling systems. **A)** Investigated taxa. Numbers in square brackets indicate the number of identified neuropeptide precursors types (out of a total of 71 types), number of identified neuropeptide GPCR types (out of a total of 41 known types) and BUSCO completeness of transcriptomes (in %). **B)** Pipeline for the identification of proneuropeptides and neuropeptide GPCRs. **C)** Scanning electron micrograph of a *Lineus longissimus* (Nemertea) larva. **D)** SEM image of a *Terebratalia transversa* (Brachiopoda) larva. **E)** SEM image of a *Phoronis muelleri* (Phoronida) larva. Scale bars: 50 µm.

To identify neuropeptide precursors, we used a combined approach of relaxed BLAST searches and pattern searches. In addition, we analysed peptide extracts of the larvae of the nemertean *Lineus longissimus* by MS/MS (see below). The resulting list of precursors was analysed and used as a new query database in a second, more stringent BLAST search to find hidden orthologs. All sequence candidates were analysed by similarity-based clustering (fig. 2) and manual curation (see Supplementary Material 1). To assess potential homologies between certain pNPs, we carried out gene structure analyses, which proved to be a powerful tool in homology assessments for neuropeptide precursors (Mair *et al.*, 2000; Yañez-Guerra *et al.*, 2020; Zhang *et al.*, 2020). With this comprehensive strategy, we identified a total of approximately 1,300 potential neuropeptide precursor candidates including different paralogs and isoforms (Supplementary Material 1, 2).

**Figure 2.**
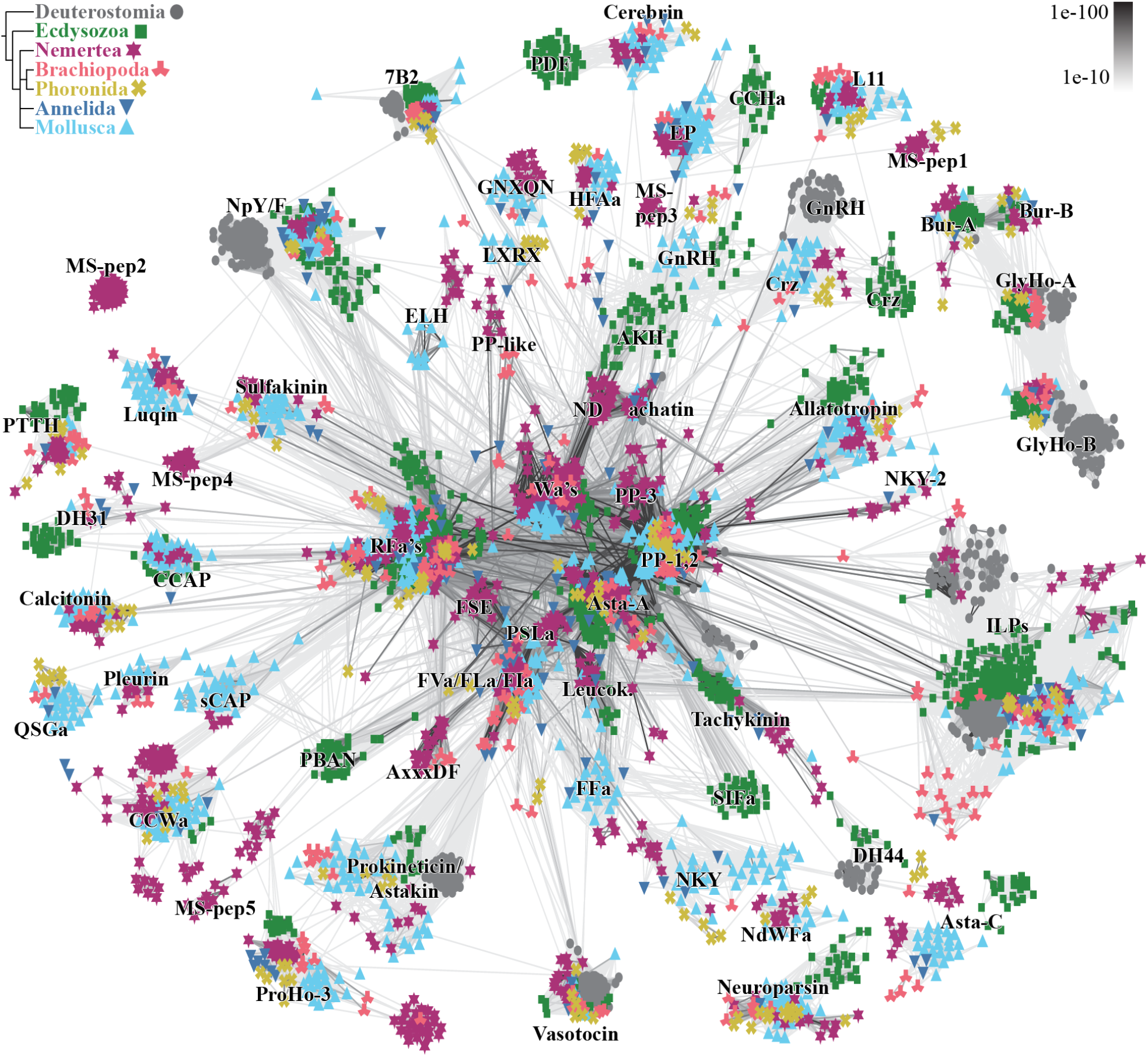
Cluster analysis of neuropeptide precursors. Connections are based on blast similarities < 1e-10 as shown on the upper right. Animal groups are colour and symbol coded as shown on the upper left. 7B2, Neuroendocrine protein 7B2; AKH, adipokinetic hormone; Alla, allatostatin; Bur, Bursicon; CCAP, crustacean cardioacceleratory peptide; CRF, corticotropin releasing factor; CRZ, corazonin, DH, diuretic hormone; ELH, egg-laying hormone; EP, excitatory peptide; ETH, ecdysis triggering hormone; GnRH, gonadotropin releasing hormone; L11, elevenin; Leucok, leucokinin; PBAN, pheromone biosynthesis activating neuropeptide; PDF, pigment dispersing factor; PTTH, Prothoracicotropic hormone; RFa’s, RFamides, Wa’s, Wamides.

To identify the full set of neuropeptide GPCRs, we used an initial HMM search (evalue 1e-10) and analysed our candidates by clustering followed by a preliminary phylogenetic analysis. We then performed a BLAST search using different bilaterian neuropeptide GPCR sequences as well as the sequences retrieved in the initial HMM search as query sequences. With this combined approach we identified over 600 neuropeptides GPCR candidates in the 23 analysed transcriptomes.

In addition to known lophotrochozoan peptidergic systems, we identified potentially novel signalling systems based on either the presence of lophotrochozoan orphan receptor groups or novel precursors. From most conserved bilaterian peptidergic signalling systems we found at least either a precursor or a receptor in one of the species of the three phyla (fig. 3, Supplementary Material 3). Only the achatin and leucokinin systems could not be detected in brachiopods and phoronids, and there was no indication of SIFamide/FFamide signalling in brachiopods. Our conclusion that achatin has been lost from phoronids and brachiopods is in contrast to a recent publication that reports a brachiopod achatin (Oliveira *et al.*, 2019). However, this was a glycine-rich non-neuropeptide sequence (lingulaAnatina.g6587.t1; XP_013387045.1) that spuriously clustered with the glycine-rich achatin pNPs.

**Figure 3.**
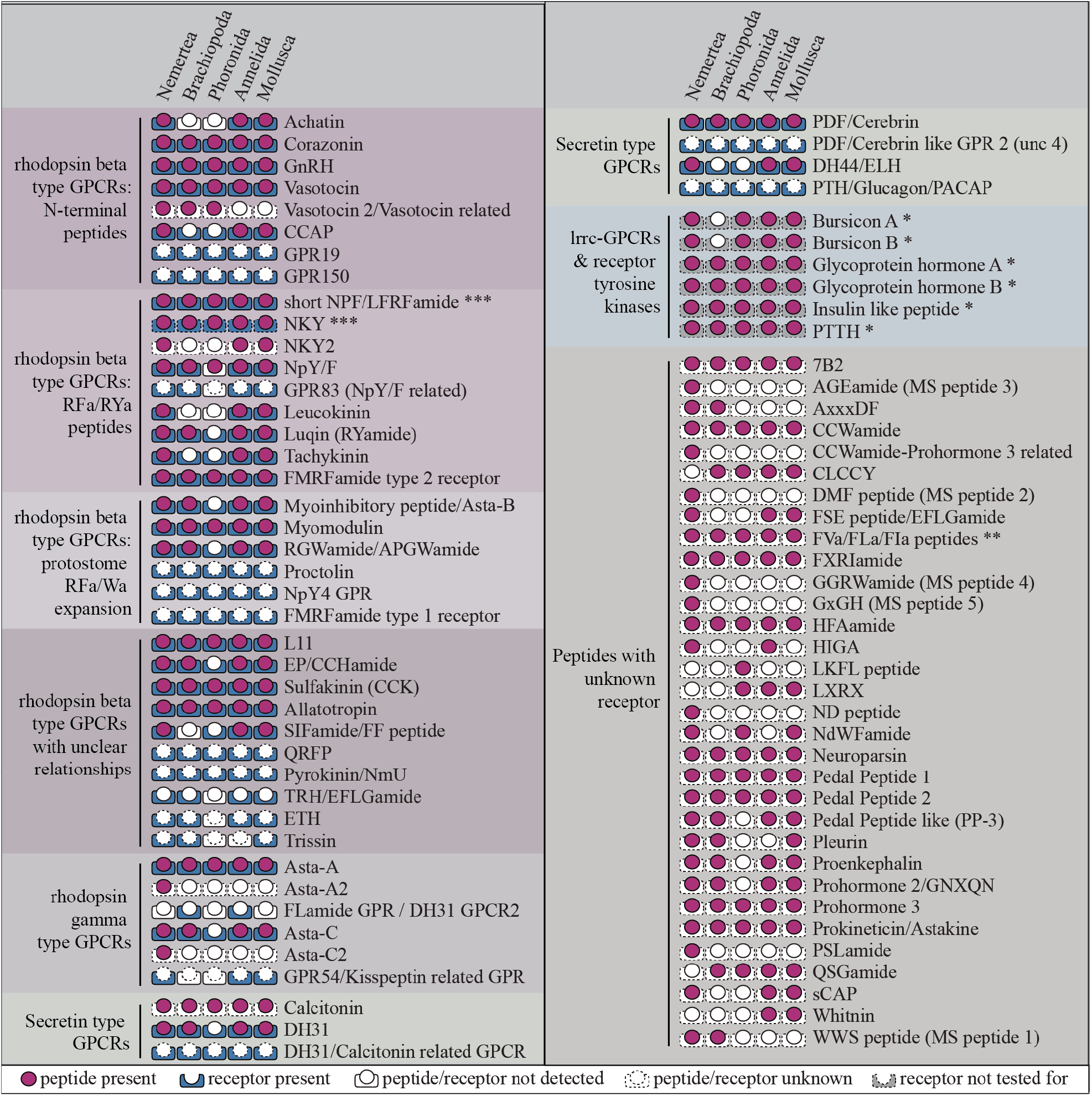
Presence of proneuropeptides and neuropeptide GPCRs. Signaling systems are sorted according to monophyletic receptor groups with consensus between this and other studies (Elphick, Mirabeau and Larhammar, 2018; Thiel *et al.*, 2018). A filled circle indicates the presence of a propeptide in at least one taxon of the corresponding clade, a filled square around the lower half of the circle indicates the presence of a receptor. A white circle or square indicates that no evidence for the presence of a precursor or receptor is present. If the circle or square has a dotted line the precursor or receptor is generally not known. * presence of leucine-rich repeat containing GPCRs or non-GPCR neuropeptide receptors was not tested for in this study. ** FVamide, FLamide and FIamides may include phylogenetically different spiralian peptides with similar C-terminal motifs. *** The depicted NKY receptors and short neuropeptide F receptors refer to the same receptor. lccr, leucine-rich repeat containing

We discovered several neuropeptide precursors that have not been identified in nemerteans, brachiopods or phoronids in a previous survey (Oliveira *et al.*, 2019). Of these previously undetected pNPs, we newly identified in nemerteans secretogranin V (7B2), calcitonin, CCWamide, CCAP, corazonin, glycoprotein hormone alpha (GlyHo alpha), GnRH, neuropeptide Y/F (NPY/F), pleurin, prohormone 2 (Proho 2) and RGWamide. Newly discovered brachiopod neuropeptide precursors include 7B2, calcitonin, CCWamide, CLCCY, corazonin, DH31, GlyHo beta, GnRH, elevenin (L11), neuroparsin, PDF/cerebrin, pleurin, ProHo 2, ProHo 3, prothoracicotropic hormone (PTTH), RGWamide, LFRFamide/short neuropeptide F (sNPF), sulfakinin and vasotocin. Newly identified precursors in phoronids include 7B2, bursicon alpha, calcitonin, corazonin, GlyHo beta, GnRH, HFAamide, L11, NPY/F, PDF/cerebrin, QSGamide and LFRFamide/sNPF.

The discovery of so many new peptidergic systems can be attributed to a large part to our thorough and iterative search strategy and the use of transcriptomes from multiple species. The full complement of identified and annotated pNP sequences, with predicted cleavage and amidation site of all investigated species is given in the Supplementary proneuropeptide list (Supplementary Material 1).

### Mass spectrometry confirms predicted cleavage sites and reveals novel pNPs

To complement our bioinformatic search, we analysed peptide extracts from larvae of the nemertean *L. longissimus* by mass spectrometry. Larvae of this species were comparatively easy to obtain in sufficient numbers, could be reared to different stages, and the reference transcriptome is of high quality (94.2% BUSCO completeness, Supplementary Material 4). Following an MS/MS run of the peptide extracts, we screened for hits against the translated transcriptome of *L. longissimus*, using a random digestion of the target database. This allowed us to randomly test for the accuracy of predicted cleavage sites and modifications, as well as to find additional, so far unknown neuropeptides.

We could confirm 46 peptides that were predicted between dibasic cleavage sites on 18 precursors which were identified by BLAST or motif search (Allatostatin A1, Allatostatin A2, Allatostatin C2, Allatotropin, ASWLDF, CCAP, CCWamide, Corazonin, DH44/ELH, FVamide/FLamide/FIamide, FMRFamide, Pedal peptide 1, Pedal peptide 2, Pedal peptide 3, Prohormone 2, PSLamide, RGWamide, SIFamide/FF peptide) (Supplementary Material 5, 6: *L. longissimus* MS precursors, *L. longissimus* MS scores). Some peptides also suggested monobasic cleavage sites, usually C-terminal to an arginine residue with a second basic amino acid residue 3-6 positions N-terminal of the cleavage site (fig. 4D, Supplementary Material 5). Such monobasic cleavage sites are known as alternative cleavage sites to the classic dibasic cleavage sites (Veenstra, 2000; Southey, Sweedler and Rodriguez-Zas, 2008), although they are less common than dibasic sites. We also detected peptides that were shorter than predicted, missing single or multiple amino acids at their C- or N-termini (Supplementary *L. longissimus* MS scores). For example, from the Pedal peptide 1 and 3 precursors we detected multiple peptides by mass spectrometry, including full length predicted peptides, as well as shorter, potentially degraded versions. Many predicted modifications like N-terminal pyroglutamination or C-terminal amidation were confirmed (if the full N-terminus or C-terminus was present).

**Figure 4.**
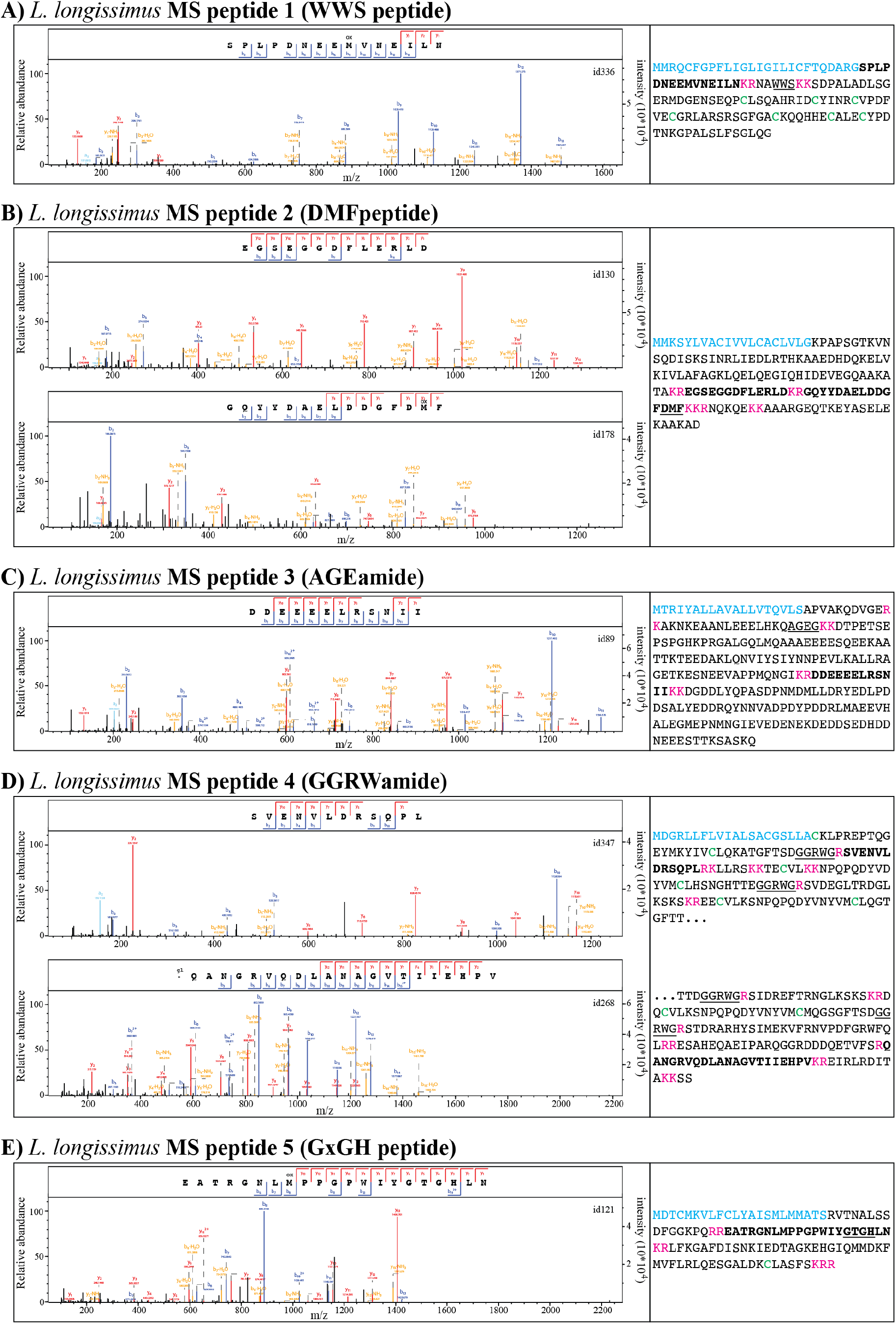
Neuropeptides discovered by mass spectrometry in *Lineus longissimus*. Spectra of novel peptides and their precursors that were identified by mass spectrometry. Spectra of peptides are shown on the left side of the panels with the corresponding precursor sequences shown to the right. Precursor sequences are marked as follows: signal peptide in blue, detected peptide in bold, name-giving sequence underlined, cleavage sites in magenta, cysteine residues in green. The precursor of MS peptide 4 (GGRWamide) is split into two partial sequences.

Finally, we identified 7 peptides derived from 5 different precursors that were not detected by BLAST or motif search (fig. 4). We added these new precursors to our BLAST screens and found homologs in other species (MS-peptides 1-5: fig. 2, Supplementary Material 1, 7). Only the WWS peptide (MS-peptide 1) precursor was also detected outside nemerteans, in phoronid species (fig. 2 MS-pep 1, Supplementary Material 1, 7), whereas the other four precursor types were only identified in nemertean species (fig. 2 MS-pep 2-5, Supplementary Material 1,7). The presence of these precursors with similar sequences between predicted cleavage sites in multiple species supports the neuropeptidergic nature of these sequences.

### The bilaterian W/Y/Famide GPCRs expanded in protostomes into multiple RF/Wamide-activated GPCRs

The phylogenetic analysis of Rhodopsin-*beta* GPCRs revealed a protostome “supergroup” of GPCRs related to RFamide and Wamide receptors (fig. 5a Tyr/Trp/Phe signaling). This protostome-specific GPCR expansion is in accordance with a previous study of xenambulacrarian GPCRs (Thiel *et al.*, 2018). The individual protostome receptor types (including myoinhibitory peptide (MIP), proctolin, RGWamide, NKY-4, FMRFamide type 1, myosuppressin) lack direct orthologs in deuterostomes. For these families, no directly orthologous propeptides have been found in deuterostomes to date (Jekely, 2013; Mirabeau and Joly, 2013). The recently reported MIP-related neuropeptides in the ambulacrarians *Saccoglossus kowalevskii* and *Apostichopus japonicus* (Zieger *et al.*, 2021) are misidentified non-neuropeptide sequences (Supplementary Material 8 Zieger et al. 2021 MIP analysis). The deuterostome receptor sequences most closely related to this supergroup are the GPR142 and GPR139 receptors from humans and *Branchiostoma floridae* (fig. 5a, Supplementary Material 9a). This further details a previously described relationship of protostome MIP (also called allatostatin-B) and proctolin receptors to the deuterostome GPR139/142 (Mirabeau and Joly, 2013).

**Figure 5.**
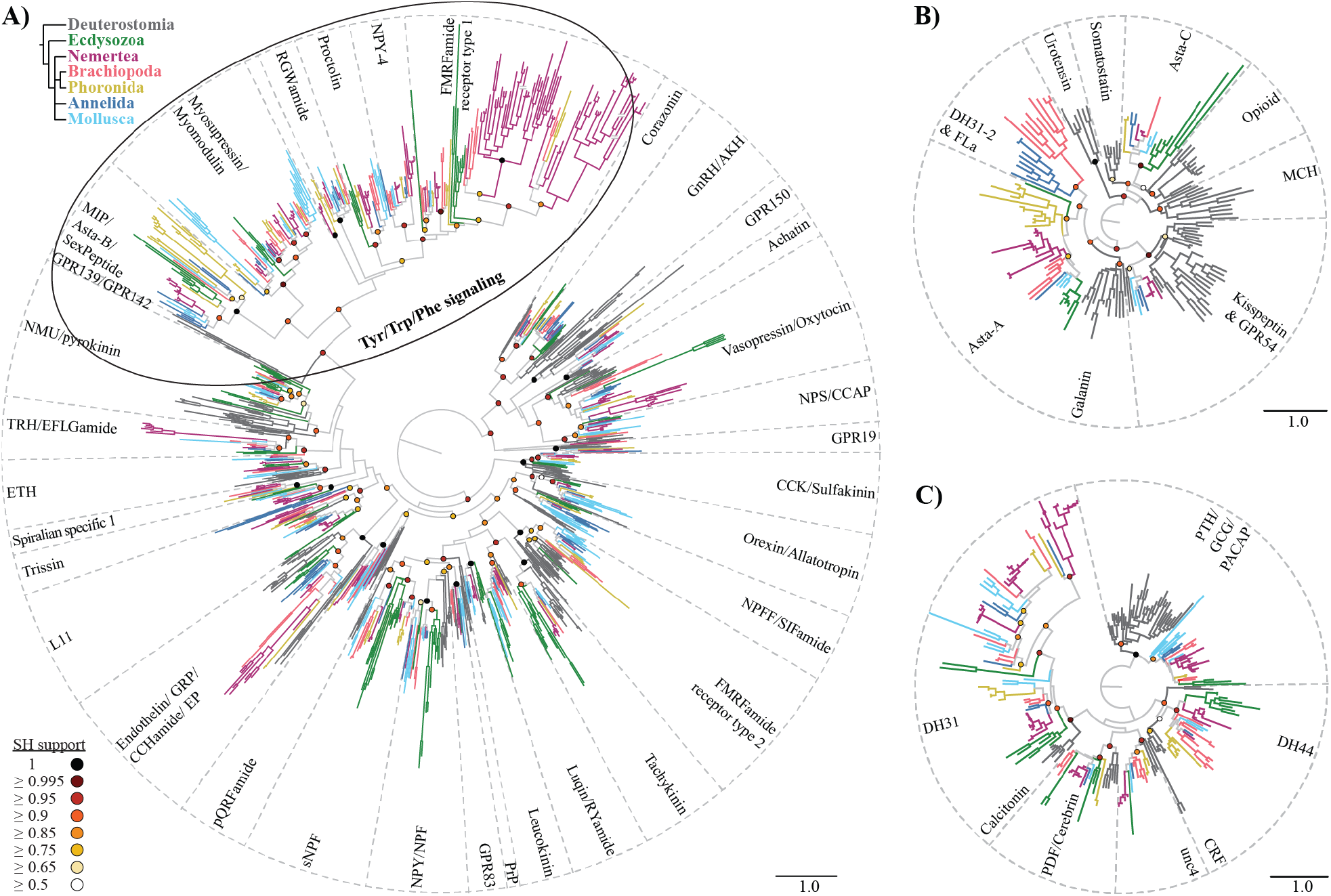
Maximum-likelihood analysis of rhodopsin and secretin-type neuropeptide GPCRs. **A)** Rhodopsin *beta* GPCRs. **B)** Rhodopsin *gamma* GPCRs. **C)** Secretin GCPRs. Terminal branches are colour-coded according to taxon as shown on the upper left. SH-aLRT support values of major nodes are colour-coded in circles as indicated on the lower left. Scale bars on the lower right of each tree indicate the inferred amino acid substitutions per site. Dashed lines demarcate orthologous receptor types. A double crossing through a branch (nemeretan FMRFamide type 1 receptors) indicates that the branch length was halfened. AKH, adipokinetic hormone; Asta, allatostatin; CCAP, crustacean cardioacceleratory peptide; CCK, cholecystokinin; CRF, corticotropin releasing factor; DH, diuretic hormone; ETH, ecdysis triggering hormone; EP, excitatory-peptide; GCG, glucagon, GHS, growth hormone secretagogue; GnRH, gonadotropin releasing hormone; GRP, gastrin releasing peptide; NMU, neuromedin-U; NPS, neuropeptide-S; NPY/NPF, neuropeptide Y, neuropeptide F; MCH, melanin concentrating hormone; MIP, myoinhibitory peptide; PACAP, pituitary adenylate cyclase-activating polypeptide; PBAN, pheromone biosynthesis activating neuropeptide; PDF, pigment dispersing factor; pQRFPa, pyroglutamylated RFamide peptide; PTH, parathyroid hormone; PRP, prolactin releasing peptide; SH, Shimodaira–Hasegawa approximate-likelihood ratio test; TRH, thyrotropin releasing hormone; VIP, vasoactive intestinal peptide.

The human GPR139 has been shown to be activated by adrenocorticotropic hormone (ACTH), α- and β-melanocyte stimulating hormone, W peptides, aromatic L- and D-amino acids, and various pharmacological antagonists with aromatic rings (Isberg *et al.*, 2014; Nøhr *et al.*, 2017; Vedel *et al.*, 2020). GPR142 is still an orphan receptor. Protostome receptors of this expanded group are activated by neuropeptides with amidated aromatic C-termini, such as MIP [Wamide], myosuppressin [RFamide], RGWamide [Wamide], *P. dumerilii* neuropeptide Y-4 [RFamide] and arthropod FMRFamide [RFamide]. Exceptions are the spiralian myomodulins and proctolin-related signalling systems. Myomodulins constitute the spiralian orthologs of the insect RFamide myosuppressin, but usually end in RLamide/RMamide and PRXamide (Oliveira *et al.*, 2019).

Our finding that most peptides that activate this supergroup of GPCRs possess an amidated aromatic amino acid on their C-terminus suggests that ancestral myomodulin-like peptides were also Famides or Wamides, similar to their arthropod orthologs. Proctolins are non-amidated peptides usually ending with a C-terminal Thr residue and are described from insects and crustaceans (Johnson *et al.*, 2003; Orchard *et al.*, 2011). The peptide that activates the orthologous spiralian proctolin receptor is so far unknown. We found members of every investigated trochozoan clade in each of the individual protostome-specific GPCR types within this supergroup. The spiralian receptors that are related to the type 1 FMRFamide receptors seem to have duplicated into at least two paralogous groups after the ecdysozoan-spiralian split, with a further subsequent expansion in the nemertean lineage. The only identified ligand of one of these type 1 FMRFamide GPCR related groups is the *P. dumerilii* NPY-4 […SRPRFamide]. The comparably high EC_50_ values (110 nM, 350 nM) in the initial NPY-4 receptor deorphanisation (Bauknecht and Jékely, 2015), however, could indicate a non specific cross-activation and the specific ligand may be a different peptide, potentially with a C-terminal RFamide motif.

As most protostome receptors in this expanded group are activated by peptides with C-terminal amidated aromatic amino acids and also the human GPR139 is activated by peptides, amino acids and other antagonists with aromatic rings, it is likely that the ancestral bilaterian GPCR was activated by peptides with C-terminal aromatic amino acids. RFamides and Wamide peptides are also present outside Bilateria, in Cnidaria and Placozoa (Walker, Papaioannou and Holden-Dye, 2009;Nikitin, 2015;Hayakawa *et al*., 2019; Koch and Grimmelikhuijzen, 2019,2020; Williams, 2020). Such non-bilaterian RFamide/Wamide peptides may be orthologous to the whole protostome expansion, rather than to specific bilaterian peptides. As this GPCR expansion happened within protostomes, it is likely that the corresponding proneuropeptides evolved and diversified in parallel with the receptors. This suggests that there are no direct orthologs of MIP, proctolin, RGWamide etc. in deuterostomes or cnidarians.

### Trochozoan CCWamides are orthologous to arthropod agatoxin-like peptides

CCWamide has so far been described in molluscs, annelids, phoronids and entoprocts. Here we identified CCWamide precursors also in nemerteans and brachiopods. We also found evidence that CCWamide is the spiralian ortholog of arthropod agatoxin-like peptides (ALPs). The U8-agatoxin peptide was first identified as a venom component in spiders (Skinner *et al.*, 1989) but ALPs have later also been found in hexapods and crustaceans (Sturm *et al.*, 2016; Christie *et* al., 2020).

A comparison of CCWamide and ALP precursors reveals a conserved precursor structure of the comparably short precursors. The signal peptide is followed by a non-conserved peptide, which is separated by a cleavage site from the C-terminal CCWamide/ALP peptide. The predicted CCWamide/ALP peptides possess eight conserved Cys residues with an aromatic amino acid often occurring between the 6th and 7th Cys and an amidated aromatic amino acid at the C-terminus (fig. 6a). A gene structure comparison of CCWamides from the mollusc *Crassostrea virginica*, the phoronid *P. australis* and the nemertean *N. geniculatus* with the ALP of the flour beetle *Tribolium castaneum* and the three agatoxin paralogues of the arachnid *Centruroides sculpturatus* reveals a conserved intron-exon structure of the precursors (fig. 6b). The precursors are usually encoded in three different exons, except for one of the three *N. geniculatus* (CCWa2) paralogs, which is encoded in five exons. All precursors encode the signal peptide in the first exon and the mature CCWamide/ALP peptide in the last exon. We also found that in the case of all the arthropod genes, one of the cleavage sites that gives rise to the mature peptide ALP is interrupted by an intron. The second to last exon contains the sequence coding for one or two basic amino acids (K/R) in the C-terminal region of the predicted protein, and the last exon contains the region coding for one basic amino acid in the N-terminal region of the predicted peptide. We found the same pattern in the *CCWamide* gene from *C. virginica*, and the *CCWamide1* and *CCWamide2* genes from *N. geniculatus*. In the case of the gene from *P. australis*, only the two basic amino acids in the C-terminal region of the second to last exon are present. Additionally, both the CCWamide and ALP genes show a conserved exon-intron junction positioned between the second and the third nucleotide of the already mentioned basic amino acid codon of the C-terminal basic amino acid. This frame is conserved in all the sequences analysed, even the CCWamide 3 from *N. geniculatus*, which does not contain such basic amino acid in the second to last exon.

**Figure 6.**
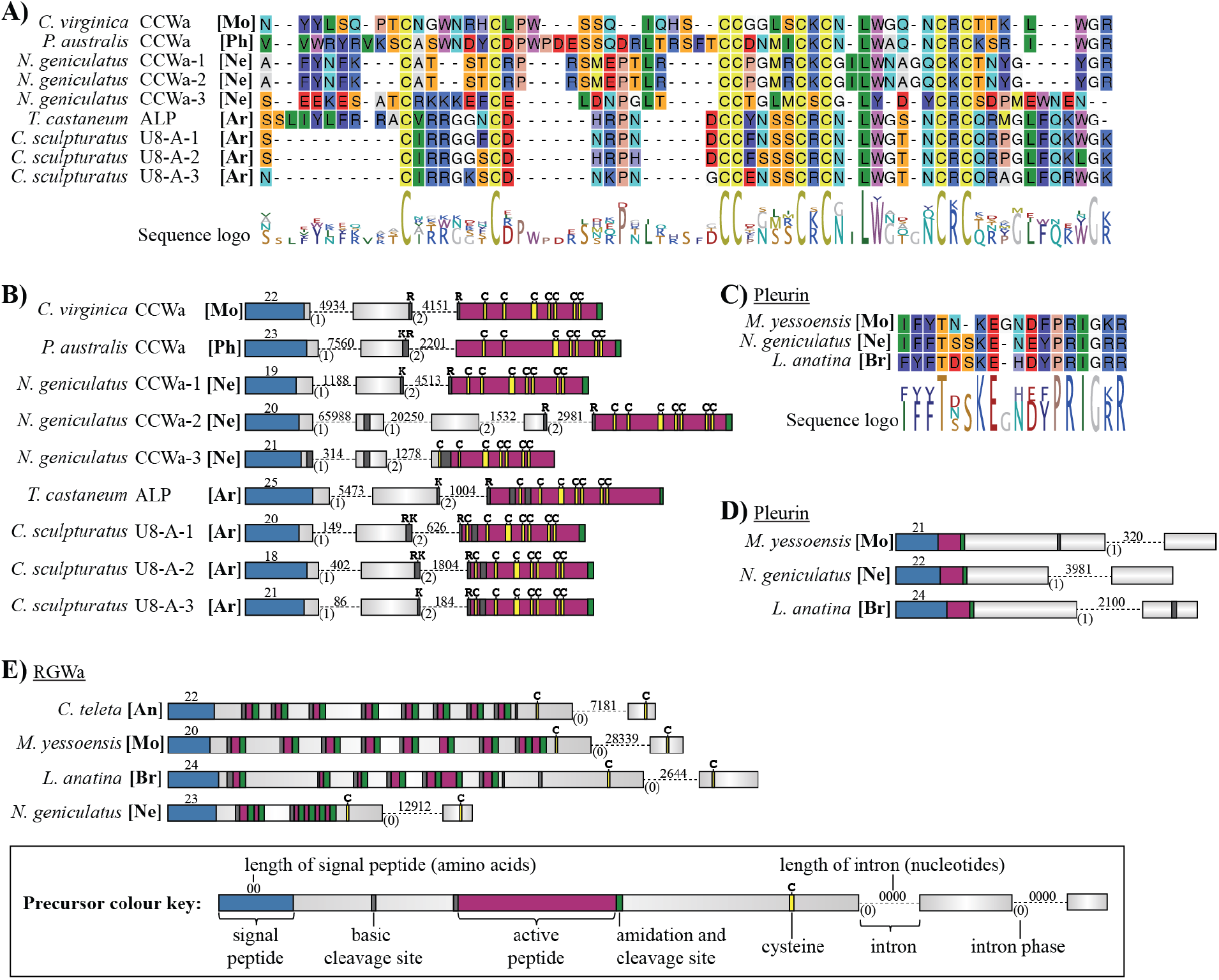
Peptide alignments and genomic precursor structure. **A)** Alignment of CCWamide and agatoxin-like peptides. **B)** Genomic exon-intron structure of CCWamide and agatoxin like peptide precursors. **C)** Alignment of pleurin peptides. **D)** Genomic exon-intron structure of pleurin precursors. **E)** Genomic exon-intron structure of APGWamide and RGWamide precursors. [An] annelid, [Ar] Arthropod, [Br] brachiopod, [Mo] mollusc, [Ne] nemertean, [Ph] phoronid.

Collectively, these findings unify the family of CCWamides from lophotrochozoans and ALP peptides from ecdysozoans, and demonstrate their single protostomian ancestry. To date, no receptor from either family has been identified. The fact that ALPs are described as part of spider venoms but orthologs are present in non-venomous arthropods and in lophotrochozoans, suggests that the neuropeptide has been recruited secondarily as a toxin. The evolution of toxin components from neuropeptides is a recurrent theme in toxin evolution and has also been reported in cone snails (Cruz *et al.*, 1987; Robinson *et al.*, 2017) and the sea anemone *Nematostella vectensis* (Sachkova *et al.*, 2020).

### Pleurin is also present outside Mollusca

Pleurin has so far been considered to be a mollusc specific neuropeptide as no orthologs have been found in other groups (Oliveira *et al.*, 2019). Many mollusc pleurin precursors encode multiple pleurin peptides in the C-terminal half of the propeptide (Veenstra, 2010; Bose *et al.*, 2017; Wang *et al.*, 2017; Oliveira *et al.*, 2019). Some bivalve species such as *Nucula tumidula* and *Mizuhopecten yessoensis*, as well as more distantly related neomeniomorph species such as *Neomenia megatrapezata* and *Wirenia argentea*, however, possess only a single peptide copy directly after the signal peptide (Zhang *et al.*, 2018). Here we identified pleurin precursors in two brachiopod and six nemertean transcriptomes. An alignment of the active peptide shows strong similarities between the peptides of the mollusc *Mizuhopecten yessoensis*, the nemertean *N. geniculatus* and the brachiopod *L. anatina* (fig. 6c). The orthology is further supported by the presence of a single genomic intron in a similar position and with the same intron phase in all three genes (fig. 6d). The presence of pleurin in nemerteans and brachiopods identifies pleurin as a trochozoan neuropeptide, which may have been lost in annelids and potentially also in phoronids. All nemertean and brachiopod pleurin precursors only have a single predicted active peptide directly after the signal peptide. The presence of only a single peptide copy in brachiopods, nemerteans and different molluscs suggests that the multi-copy nature of various mollusc pleurins evolved from an ancestral single-copy pleurin precursor.

### APGWamide is the mollusc ortholog of RGWamide

Precursors encoding the RGWamide peptides have been described from different annelid species (Martín-Durán *et al.*, 2020; Veenstra, 2011; Conzelmann, Williams, Krug, *et al.*, 2013) and the RGWamide receptor was identified in the annelid *P. dumerilii (Bauknecht and Jékely, 2015)*. A previous study proposed that the annelid RGWamide may be the ortholog of the mollusc APGWamide (Veenstra, 2011), but no further explanation was given and APGWamide was classified as a mollusc specific peptide (Oliveira *et al.*, 2019). Here, we identified RGWamide precursors in nemerteans and brachiopods, and orthologs of the *P. dumerilii* RGWamide receptor in species of all five trochozoan clades (fig. 2, fig. 5a). The receptor analysis reveals that the RGWamide signalling system is generally present in all trochozoans, including molluscs, and that it is part of the protostome specific RFamide/Wamide signalling system expansion. An investigation of the precursors and mature peptides of RGWamides and APGWamides reveals several similarities which support the orthology of these neuropeptides. Most APGWamide and RGWamide precursors contain multiple peptide copies in their more N-terminal region and two cysteine residues at their C-terminus (fig. 6e, Supplementary Material 7). The two cysteine residues are separated by 20-30 amino acids without any RGWamide/APGWamide copy in between. The gene structure analysis of the APGWamide precursor of *Mizuhopecten yessoensis* (Mollusca) and the RGWamide precursors of *Capitella teleta* (Annelida), *L. anatina* (Brachiopoda) and *N. geniculatus* (Nemertea) reveals a single phase-0 intron between the exons coding for the two C-terminal Cys residues (fig. 6e). Furthermore, precursors of heteronemertean RGWamides also contain the tetrapeptide APGWamide as their most N-terminal predicted active peptide. In addition, several of the brachiopod and nemertean precursors encode active peptides that follow each other in direct succession, leading potentially to a cleavage into GWamide dipeptides instead of RGWamide. Such GWamide dipeptides have also been predicted in the APGWamide precursor of the mollusc *Lottia gigantea* (Veenstra, 2010) *and have been identified by mass spectroscopy in Sepia officinalis* as a product of alternative processing (Henry and Zatylny, 2002). Overall, our findings demonstrate that molluscs APGWamides and RGWamides from other spiralians are orthologous. We therefore suggest that the molluscan RGWamide GPCR orthologs that we used in our phylogenetic analysis (*Lottia gigantea*: XP_009046545, XP_009052844; *Crassostrea gigas*: EKC25208; Supplementary Material 9) are likely to be activated by molluscan APGWamide peptides.

### Vasotocin duplications in spiralians and diversification in nemerteans

Vasotocin orthologs can be found throughout bilaterians. Within the vertebrate lineage, the whole signalling system (pNPs and GPCRs) expanded and led to the evolution of the paralogous vasopressin and oxytocin systems. The nomenclature of this peptide is confusing with different names across animals including cephalotocin in cephalopods (Bardou *et al.*, 2010), asterotocin in the starfish *Asterias rubens* (Semmens *et al.*, 2016), inotocin in insects (Stafflinger *et al.*, 2008) and vasotocin, inotocin or mesotocin for different teleost homologs (Van den Dungen *et al.*, 1982; Larson, O’Malley and Melloni, 2006). Here we use vasotocin derived from combining the names of the mammalian paralogs vasopressin/oxytocin. The peptide precursor of vasotocin related neuropeptides is strongly conserved throughout bilaterians: the N-terminal signal peptide is directly followed by a single copy of the bio-active vasotocin peptide, which is then followed by a large neurophysin domain that is characterised by conserved Cys residues (fig. 7a). The precursor is therefore often referred to as a combination of the active peptide (vasotocin, oxytocin,…) and neurophysin (e.g. vasotocin-neurophysin). In our search we detected two vasotocin paralogs in the linguliform brachiopods *L. anatina* and *G. pyramidata*, and the craniiform brachiopod *N. anomala*, but only a single ortholog in the rynchonelliform brachiopods. We also identified two vasotocin paralogs in the phoronid species *P. vancouverensis* and *P. australis*, as well as in the paleonemertean *T. polymorphus*. In hetero- and hoplonemerteans, however, we detected a novel peptide, which is potentially related but strongly diverged. Similar to other vasotocin-like peptides, the precursor of this novel peptide encodes a single copy of the active peptide, which is then followed by a C-terminal neurophysin domain. The predicted novel peptides, however, lack the strongly conserved cysteine residues found in vasotocin-like peptides and possess an N-terminal glutamine instead (fig. 7b). The active peptides of hoplonemerteans are potentially amidated at their C-terminus, similar to the C-terminal amidation of classical vasotocin related peptides. The active peptides of the heteronemertean species, however, seem to have a non-amidated C-terminus. The presence of a neurophysin-like domain in a non-vasotocin precursor has also been described for echinoderm NG peptides (Semmens *et al.*, 2015). The neurophysin domain in NG peptides, however, dates back to a far more ancestral bilaterian duplication that subsequently led to the evolution of recent vasotocin-related neuropeptides from one paralog and neuropeptide S/NG peptide/CCAP from the other paralogous system (Semmens *et al.*, 2015; Semmens and Elphick, 2017).

**Figure 7.**
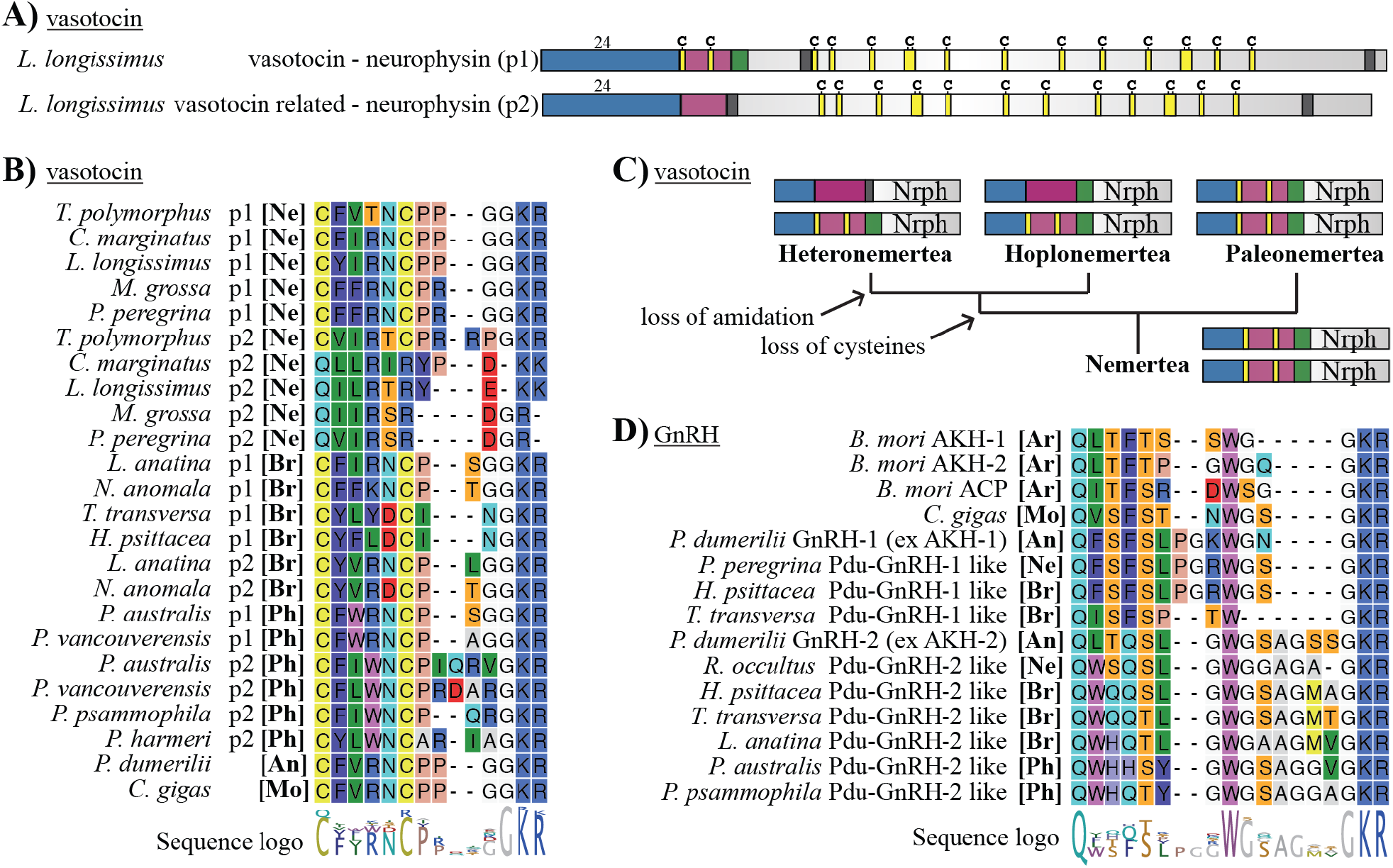
Vasotocin and GnRH related peptides. **A)** Precursor structure of *L. longissimus* vasotocin paralogs. Length of signal peptide in amino acids is shown above the precursors, rest is to scale. **B)** Alignment of vasotocin and related peptides. **C)** Evolution of vasotocin-related proneuropeptides in nemerteans. Precursors are not to scale. **D)** Alignment of GnRH and related peptides. *B. mori Bombyx mori, C. gigas Crassostrea gigas*, Nrph neurophysin, p1 paralog 1, p2 paralog 2, Pdu *Platynereis dumerilii*, [An] annelid, [Ar] Arthropod, [Br] brachiopod, [Mo] mollusc, [Ne] nemertean, [Ph] phoronid.

To assess the evolutionary scenarios that led to the vasotocin-related paralogs in nemerteans, brachiopods and phoronids, we calculated two maximum likelihood trees (Supplementary Material 10). One tree is based on the alignment of the whole precursors, excluding the signal peptide and the other one is based on the neurophysin domain only. Both phylogenies suggest that the strongly diverged hoplo- and heteronemertean peptides are related to the vasotocin paralog 2 of the paleonemertean *T. polyphemus*, which constitutes a “classic” vasotocin paralog (fig. 7b). This suggests that the novel hoplo- and heteronemertean peptides are strongly divergent vasotocin paralogs with a stepwise loss of the two characteristic cysteine residues in the ancestral lineage leading to Hoplonemertea and Heteronemertea and a further loss of the amidation site in the heteronemertean lineage (fig. 7c). The evolution of the brachiopod and phoronid vasotocin paralogs is less clear. Both trees suggest a common origin of the phoronid paralog 2 and brachiopod paralog 2 precursors (supplementary Material 10), which is the precursor that was not detected in rhynchonelliform brachiopods. The neurophysin tree further suggests a common origin of the phoronid paralog 1 and brachiopod paralog 1, suggesting that the phoronid and brachiopod paralogs are derived from a single duplication event. This is in contrast to the vasotocin-neurophysin tree, which is unclear about the origin of the linguliform brachiopod paralog 1 precursors in relation to other trochozoan species. There is no support for a single duplication event in the ancestral trochozoan lineage, as annelid and mollusc species generally seem to possess only a single ortholog (Martín-Durán *et al.*, 2020; Veenstra, 2010, 2011; Conzelmann, Williams, Krug, *et al.*, 2013; Stewart *et al.*, 2014). We did not identify vasotocin GPCR duplications that generally correspond to the presence of two vasotocin paralogs in the investigated clades. The only duplication of vasotocin-related GPCRs that we detected was in nemerteans – specifically in several heteronemertean species and in the hoplonemertean *Nipponemertes* sp (Supplementary Material 9a).

### Ancestral duplication of GnRH related neuropeptides in spiralians

GnRH-like neuropeptides and their receptors underwent several duplications in different bilaterian lineages.The protostome-deuterostome last common ancestor already possessed two paralogous GnRH-like signalling systems called GnRH and corazonin (Tian *et al.*, 2016; Zandawala, Tian and Elphick, 2018). Along the arthropod lineage, the GnRH system duplicated to give rise to the paralogous AKH and ACP systems (Hansen *et al.*, 2010; Hauser and Grimmelikhuijzen, 2014; Tian *et al.*, 2016; Zandawala, Tian and Elphick, 2018), while the corazonin system has been maintained. In addition to these ancestral duplications, more recent duplications of GnRH or corazonin precursors occurred in some annelids (Martín-Durán *et al.*, 2020; Veenstra, 2011; Conzelmann, Williams, Krug, *et al.*, 2013).

We could find both corazonin and GnRH precursors and their putative GPCRs in nemerteans, brachiopods and phoronids (fig. 5a, fig. 7d, Supplementary Material 1). In addition, we found evidence for a duplication of GnRH neuropeptides in the ancestral lophotrochozoan lineage.

There has been some confusion over the annotation of the GnRH and corazonin precursors in many trochozoan species in the past. This was recently clarified based on peptide sequence similarity (Zandawala, Tian and Elphick, 2018) and receptor deorphanisation in the annelid *Platynereis dumerilii* (Williams *et al.*, 2017; Andreatta *et al.*, 2020), although there is still some confusion left due to nonspecific receptor activation (Andreatta *et al.*, 2020). In short, there have been two GnRH like paralogs described in *P. dumerilii*, originally named AKH-1 and AKH-2 (Conzelmann, Williams, Krug, *et al.*, 2013). AKH-1 was renamed to GnRH-1 based on sequence similarity of the peptide (Zandawala, Tian and Elphick, 2018). A deorphanisation assay showed, however, that this GnRH1 peptide does not activate any of the two *P. dumerilii* GnRH GPCRs, but at high concentrations the two *Platynereis dumerilii* corazonin receptors and was therefore referred to as GnRHL3. GnRH2 specifically activated the two *Platynereis dumerilii* GnRH receptors (Andreatta *et al.*, 2020). The two peptides that have originally been described as GnRH-1 and GnRH-2 (Conzelmann, Williams, Krug, *et al.*, 2013) have been renamed to Corazonin-1 and Corazonin-2 based on sequence similarity as well as receptor activation (Williams *et al.*, 2017; Andreatta *et al.*, 2020). The predicted *P. dumerilii* GnRH2 peptide has a longer C-terminus than other GnRH-like peptides and to our knowledge no similar paralog has been described in any mollusc species so far. In the nemertean *R. occultus*, the brachiopods *T. transversa, H. psittacea* and *L. anatina*, as well as in the three *Phoronis* species we found peptides that are similar to the *Platynereis dumerilii* GnRH-2 with its extended C-terminus. In the nemertean *P. peregrina* and the brachiopods *H. psittacea* and *T. transversa*, we found peptides that are nearly identical to the *P. dumerilii* GnRH-1 and the GnRH that is known from molluscs (fig. 7d). The presence of both paralogs in the annelid, nemertean and brachiopod lineage and the lack of GnRH type 1 in phoronids and GnRH type 2 in molluscs indicates a duplication of the GnRH neuropeptide precursor in the ancestral trochozoan lineage. This duplication was then potentially followed by a secondary loss of GnRH type 2 in the mollusc lineage and GnRH type 1 in the phoronid lineage, although we cannot rule out that the lack of GnRH type 1 in phoronids is due to a lack of detection.

We could not detect any ancestral duplications of GnRH-related GPCRs. The GnRH GPCR branch is divided into two main branches in our analysis, but only sequences of the annelids *P. dumerilii* and *Capitella teleta* are represented in both branches, while brachiopod, phoronid, nemertean and mollusc sequences are either present in one or the other branch. There may be a second GnRH receptor in annelids, as only the GnRH-2 but not the shorter GnRH-1 peptide activates the canonical GnRH receptor in *P. dumerilii* (Andreatta *et al.*, 2020). *In contrast, Crassostrea gigas* GnRH, which is more similar to the *P. dumerilii* GnRH-1, activated the typical GnRH receptor (Li *et al.*, 2016).

## Discussion

In this study, we analyse and describe the neuropeptide and neuropeptide GPCR complement of nemerteans, brachiopods and phoronids. Our study demonstrates how an increased taxon sampling and a comprehensive search strategy can help identify peptidergic systems and unravel their evolution. The inclusion of multiple taxa helped to identify the complement of larger taxonomic groups since even species with high quality transcriptomes (Supplementary Material 4) do not have the full set of neuropeptides or neuropeptide GPCRs in the dataset (Supplementary Material 3). The use of multiple search approaches, followed by the use of the initial results as new search queries helped to increase the discovery of pNPs and neuropeptide GPCR complement within species. The combination of sequence alignments and gene structure analyses helped to determine the homologies of CCWamides and ALPs, as well as APGWamides and RGWamides. Using our combined approach it was also possible to identify pleurin as an ancestral spiralian neuropeptide. Our dataset of neuropeptide precursors (Supplementary Material 1,2) and neuropeptide GPCRs (Supplementary 9 in combination with Supplementary Material 11-13) can be used as a resource when identifying signalling systems in unexamined taxa or when trying to identify new peptide-receptor pairs.

There are several outstanding questions about neuropeptide-receptor systems in protostomes (see also fig. 3). The receptor for CCWamides and their ecdysozoan ALP orthologs is currently unknown. This receptor is expected to be present in both ecdysozoans and lophotrochozoans and may have undergone an expansion in nemerteans, in parallel with the expansion of CCWamide pNPs (Fig. 2, Supplementary Material 1). Another protostome pNP without a known receptor is prohormone 3. We detected a second prohormone 3 in nemerteans and the brachiopod *Laqueus californicus* (Fig. 2, Supplementary Material 1) and a potential third paralog in nemerteans with similarities to CCWamide (Fig. 2, Supplementary Material 1). Prokineticin/astakine-related pNPs also have no known receptor, which is exceptional, as the deuterostome prokineticin GPCRs seem to be ancestral to bilaterians but have been lost in protostomes (Mirabeau and Joly, 2013; Thiel *et al.*, 2018). It may be that some of these peptides activate receptors outside the rhodopsin or secretin GPCR family. Potential and poorly explored receptor families include peptide gated ion channels (Golubovic *et al.*, 2007; Schmidt *et al.*, 2018) and leucine-rich repeat-containing GPCRs like those for glycoprotein hormone or relaxin related signaling (Jekely, 2013; Roch and Sherwood, 2014).

In addition to neuropeptides with unknown receptors, there are also lophotrochozoan orphan GPCRs for some of which only the deuterostome or ecdysozoan ortholog has been deorphanised. These include orthologous GPCRs of trissin receptors, kisspeptin receptors, PACAP/PTH receptors and proctolin receptors. The molluscan peptide PKYMDT has been proposed to be a potential ortholog of the insect neuropeptide proctolin (Veenstra, 2010) but an orthologous peptide has so far not been identified in any other lophotrochozoan. The endogenous ligands of lophotrochozoan FMRFamide type-1 receptors and ecdysozoan type-2 receptors are also unknown. The FMRFamide type-1 receptor was deorphanised from *Drosophila melanogaster* and is activated by different FMRFamide-like peptides with the highest sensitivity to endogenous *Drosophila* FMRFamide peptides (Cazzamali and Grimmelikhuijzen, 2002; Meeusen *et al.*, 2002). In lophotrochozoans, however, FMRFamide activates a completely different receptor (Bauknecht and Jékely, 2015; Thiel *et al.*, 2017), the FMRFamide type-2 receptor. The corresponding lophotrochozoan ligand of the FMRFamide type-1 receptor is unknown. As these receptors are part of the protostome W/Y/Famide activated GPCR expansion, it is likely that the activating peptide has an amidated aromatic amino acid at its C-terminus.

Another uncertainty relates to the receptors of NKY and sNPF peptides in lophotrochozoan. A large-scale screen identified a receptor for the two *P. dumerilii* NKY peptides (Bauknecht and Jékely, 2015). In our phylogeny, this GPCR is orthologous to deorphanised mollusc, arthropod and nematode sNPF receptors (Mertens *et al.*, 2002; Garczynski, Crim and Brown, 2007; Bigot *et al.*, 2014; Christ *et al.*, 2018; Matsumoto *et al.*, 2019). This grouping is consistent with a recent phylogenetic analysis of NPY/NPF and sNPF/PrRP receptors (Yañez-Guerra *et al.*, 2020). Both NKY and sNPF peptides have an RY/Famide motif, albeit NKY peptides are very long (up to 43 residues compared to five-six in sNPF). However, the *P. dumerilii* NKY receptor can also be activated in the high nanomolar range by FMRFamide (Bauknecht and Jékely, 2015). Likewise, the *C. gigas* NKY and FMRFamide peptides cross-activate the *C. gigas* sNPF receptor (Yañez-Guerra *et al.*, 2019), also in the high nanomolar range. *P. dumerilii* sNPF peptides (originally annotated as RYamide (Conzelmann, Williams, Krug, *et al.*, 2013)) have not yet been tested on the *P. dumerilii* NKY receptor and may represent the endogenous ligands (Yañez-Guerra *et al.*, 2019). If this turns out to be the case, this would suggest that NKY peptides have a different endogenous receptor.

Spiralian phylogeny and nomenclature has been in a flux with the clades collectively referred to as Spiralia (Hejnol *et al.*, 2009; Edgecombe *et al.*, 2011; Dunn *et al.*, 2014) or Lophotrochozoa (Telford, Budd and Philippe, 2015; Kocot *et al.*, 2017; Laumer *et al.*, 2017; Bleidorn, 2019; Marlétaz *et al.*, 2019). Some trees suggest a close relationship of molluscs, annelids, nemerteans, brachiopods and phoronids (referred to as Trochozoa). In other analyses, however, Trochozoa are paraphyletic with gastrotrichs, entoprocts, ectoprocts or platyhelminthes more closely related to the individual trochozoan clades. Regardless of the exact phylogeny, with the here presented evidence from nemerteans, brachiopods and phoronids in combination with previous studies on annelids and molluscs, we have now a well-sampled complement of neuropeptide signaling systems in these major lineages. Some spiralian branches still lack deeper sampling, including Syndermata, Gnathostomulida and Micrognathozoa (Gnathifera) (Dunn *et al.*, 2014; Laumer *et al.*, 2017) as well as Chaetognatha (Bleidorn, 2019; Marlétaz *et al.*, 2019). One study that included three species of bdelloid rotifers from the genus *Rotaria* only identified very few pNPs (Oliveira *et al.*, 2019). An in-depth survey in gnathiferans with an increased taxon sampling is the next frontier in the comparative genomics of spiralian peptidergic systems and could help to clarify some of the remaining uncertainties.

## Material and Methods

### Transcriptomic resources

For increased taxon sampling, we collected transcriptomes of different nemertean, brachiopod and phoronid species. Transcriptomes of the phoronid *Phoronis australis* (downloaded 23rd March 2020), the brachiopod *Lingula anatina* (downloaded 23rd March 2020) and the nemertean *Notospermus geniculatus* (downloaded spring 2019) were retrieved from https://marinegenomics.oist.jp (see also (Luo *et al.*, 2018)). Transcriptomes of the phoronids *Phoronis psammophila* and *Phoronis vancouverensis*, the brachiopods *Glottia pyramidata, Hemithris psittacea, Laqueus californicus* and *Novocrania anomala*, the nemerteans *Cephalothrix hongkongiensis, Cerebratulus marginatus, Lineus lacteus, Malacobdella grossa, Paranemertes peregrina* and *Tubulanus polymorphus* were downloaded from (10.5061/dryad.30k4v), a public dataset made available by (Kocot *et al.*, 2017)). Transcriptomes of the phoronid *Phoronopsis harmeri*, the brachiopods *Novocrania anomala* and *Terebratalia transversa*, and the nemerteans *Lineus longissimus* and *Lineus ruber* were assembled according to (Cannon *et al.*, 2016). Sequencing data of the nemerteans *Baseodiscus unicolor* (SRR1505175), *Cerebratulus* spec. (SRR1797867), *Nipponemertes* spec. (SRR1508368) and *Riseriellus occultus* (SRR1505179) were retrieved from NCBI, trimmed with Trimmomatic 0.35 (Bolger, Lohse and Usadel, 2014), error-corrected with SPAdes 3.6.2 (Nurk *et al.*, 2013) and assembled with Trinity 2.2.0 (Grabherr *et al.*, 2011). The two *N. anomala* transcriptomes were merged and sequence redundancy was reduced in all final transcriptomes with CDhit-EST (Li and Godzik, 2006; Fu *et al.*, 2012) (using a threshold of 95% similarity). Transcriptomes were translated into protein sequences with TransDecoder (TransDecoder, no date) http://transdecoder.github.io/ and a defined minimum length of 60 amino acids. A completeness-assessment of each transcriptome was performed with BUSCO v4.0.6 (Seppey, Manni and Zdobnov, 2019) using the protein mode and the lineage dataset metazoa_odb10 (Creation date of the database: 20-11-2019, number of BUSCOs: 954).

Depicted relationships of nemertean species are based on (Andrade *et al.*, 2014; Kvist, Chernyshev and Giribet, 2015), the relationships of phoronids are based on (Santagata and Cohen, 2009), and the relationship of brachiopod species is based on (Kocot *et al.*, 2017;

Marlétaz *et al.*, 2019), and a taxonomic classification according to http://www.marinespecies.org/ (status: July 2020).

### Identification and analysis of neuropeptide GPCRs

To identify potential neuropeptide receptors, we analysed the translated nemertean, brachiopod and phoronid transcriptomes for neuropeptide GPCRs. We did not include leucine-rich repeat containing GPCRs such as relaxin or glycoprotein hormone-related GPCRs, or non-GPCR neuropeptide receptors such as the insulin or PTTH receptor tyrosine kinases.

Multiple sequence alignments of Rhodopsin GPCR A (PF00001) and Secretin GPCR B (PF00002) were downloaded from the PFAM database (https://pfam.xfam.org). The sequence alignments were used for a Hidden Markov Model (HMM) search using hmmer-3.1b2 (Eddy, 2011) with an e-value cutoff of 1e-10. The resulting sequences were analysed using Phobius (Käll, Krogh and Sonnhammer, 2007) to predict the number of transmembrane domains and only sequences with a minimum of 4 and maximum of 8 transmembrane domains were kept. For our further analysis we used previously analysed neuropeptide GPCRs as reference sequences (Mirabeau and Joly, 2013; Bauknecht and Jékely, 2015; Thiel *et al.*, 2018; Yañez-Guerra *et al.*, 2020), complemented with sequences that were retrieved from NCBI for receptor types that were initially underrepresented. To determine whether the candidate sequences are indeed potential neuropeptide receptors we used CLANS (Frickey and Lupas, 2004) in initial cluster analysis. The sequence candidates were then separated into rhodopsin beta, rhodopsin gamma and secretin neuropeptide GPCRs for further phylogenetic analyses. Sequence candidates were aligned with MAFFT version 7, using the iterative refinement method E-INS-i and the standard scoring matrix BLOSUM62 (Katoh and Standley, 2013). Alignments were trimmed with TrimAl in automated mode (Capella-Gutiérrez, Silla-Martínez and Gabaldón, 2009). Maximum-likelihood trees were calculated with the online application of PhyML 3.0 (Guindon *et al.*, 2010) (http://www.atgc-montpellier.fr/phyml/) using the model LG+G4 which was automatically selected by the Smart Model Selection tool (Lefort, Longueville and Gascuel, 2017) and aLRT-SH-like branch support with 1000 replicates. Based on these trees we determined robust groups of neuropeptide GPCR candidates. We then used this set of neuropeptide GPCR sequences as reference sequences for a second search in all translated transcriptomes using this time a BLASTp search (1e-50). To include potentially new receptor types or paralogs that might have been missing in our reference database, we also included transcriptomes of *Crassostrea gigas* and *Daphnia pulex* in this second search (downloaded from http://metazoa.ensembl.org, 05.05.2020). All new candidates from the BLAST search were added to the existing list and a second phylogenetic analysis was carried out using the same methodology as before.

### Identification and analysis of neuropeptide precursors

To identify neuropeptide precursor sequences we combined a relaxed tBLASTn search (1e-1) with a pattern search and then used the combined results as new reference sequences in a second, more stringent tBLASTn search (1e-5). We compiled a reference database of neuropeptide precursor sequences from previously analysed bilaterian datasets (Martín-Durán *et al.*, 2020; Conzelmann, Williams, Krug, *et al.*, 2013; Jekely, 2013; Adamson *et al.*, 2015; Bose *et al.*, 2017; Oliveira *et al.*, 2019), partially complemented with sequences retrieved from ncbi (Supplementary Material 25). For the pattern search, we scanned the translated transcriptomes for potential neuropeptide precursors that code for multiple peptide copies using regular expressions. To reduce a high number of false-positive hits in the pattern search, we first sorted for sequences that possess an N-terminal signal peptide with the command line version of SignalP 4.1 (Petersen *et al.*, 2011). We then used Unix bash regex commands to scan these secretomes for sequences with at least three repeats of amidation sites that are separated by 2-25 amino acids or at least five repeats of dibasic cleavage sites that are separated by 5-25 amino acids (grep -E -B 1 -e “((.{2,25}G[KR][KR])|(.{2,25}[KR](.{1}|.{3}|.{5})G[KR*])){3,}” -e “(.{5,25}[KR][KR]){5,}” “input_transcriptome.fasta” | grep -E -v “[-][-]” > output_file.fasta). The hits of the multicopy peptides from the BLAST and regex search were combined for each species, sequence redundancy was reduced using CDhit (threshold 0.95) and all candidates longer than 650 amino acids were deleted. The resulting sequence collection was clustered together with the reference sequences using CLANS (Threshold 1e-5). Non-connected sequences were deleted and samples from every cluster were taken, blasted on NCBI and checked for the existence of a signal peptide. The general precursor structure was then manually curated based on the presence of basic cleavage sites, amidation sites and conserved cysteines. All detected nemertean, brachiopod and phoronid neuropeptide precursors were then used as new reference sequences in a second tBLASTn search (Threshold 1e-5) to detect potential hidden orthologs. New sequences were added to the existing collection and the precursors were checked again. The final groups of sequences were manually annotated based on precursor structure, peptide motifs and similarity to known neuropeptides. Analysis of the exon-intron structure of genes encoding different neuropeptide precursors was performed using the SPLIGN online alignment tool (Kapustin *et al.*, 2008).

### Peptide extraction and mass spectrometry (LC-MS/MS)

Peptide extraction and mass spectrometry was done as previously described (Thiel *et al.*, 2018) with slight modifications. We collected about 100 lab-spawned larvae of the nemertean *L. longissimus* in different stages. Specimens were starved for 1 day and rinsed several times with sterile seawater. Larvae were collected into an Eppendorf cup and centrifuged for 30 seconds in a table-top centrifuge. The seawater was replaced with ultrapure water, larvae were immediately centrifuged again, and the water was replaced with an extraction buffer (90% Methanol, 9% acetic acid, 1% distilled water). The mixture was ground with a pestle and vortexed vigorously. The suspension was centrifuged at about 13,000 x g for 20 minutes at 4°C. The supernatant was collected, evaporated in a vacuum concentrator and dissolved in 300 µl ultrapure water. Neuropeptide mixtures were reduced and alkylated as described in (Borchert *et al.*, 2010) and desalted with C18 StageTips (Rappsilber, Mann and Ishihama, 2007).

LC-MS analysis was carried out on an EasyLC nano-UHPLC coupled to a Q Exactive HF mass spectrometer (both Thermo Fisher Scientific). Separations of the peptide mixture was done as previously described (Kliza *et al.*, 2017) with slight modifications. Peptides were eluted with an 87-min segmented gradient of 10-33-50-90% HPLC solvent B (80% acetonitrile in 0.1% formic acid). The mass spectrometer was operated in the positive ion mode. Full scan was acquired in the mass range from m/z 300 to 1650 at a resolution of 120,000 followed by HCD fragmentation of the seven most intense precursor ions. High-resolution HCD MS/MS spectra were acquired with a resolution of 60,000. The target values for the MS scan and MS/MS fragmentation were 3×10^6^ and 10^5^ charges with a maximum fill time of 25 ms and 110 ms, respectively. Precursor ions were excluded from sequencing for 30 s after MS/MS. MS/MS on singly charged precursor ions was enabled. The acquired MS raw files were processed using the MaxQuant software suite v.1.5.2.8 (Cox and Mann, 2008).

Extracted peak lists were submitted to database search using the Andromeda search engine (Cox *et al.*, 2011) to query target-decoy databases consisting of the translated Lineus longissimus transcriptome sequences and commonly observed contaminants (285 entries). No enzyme specificity was defined. The minimal peptide length was set to four amino acids. The initial precursor mass tolerance was set to 4.5 ppm, for fragment ions a mass tolerance of 20 ppm was used. Carbamidomethylation of cysteines was defined as fixed modification in the database search. A set of expected variable modifications was defined in the database search: pyroglutamate formation of N-terminal glutamine, oxidation of methionine, acetylation of the peptide N-terminus, amidation of the peptide C-terminus, and sulfation of tyrosine. False discovery rates were set to 1% at peptide, modification site, and protein group level, estimated by the target/decoy approach (Elias and Gygi, 2007). The data are available via ProteomeXchange with identifier PXD023147.

## Supplementary Material

### The following supplementary files are directly provided with this article

Supplementary Material 1: Full list of all identified proneuropeptides with highlighted cleavage sites, signal peptides and other characteristics.

Supplementary Material 2: Full list of all identified proneuropeptides in fasta format. Supplementary Material 3: Table with types and number of identified pNPs and GPCRs for each species.

Supplementary Material 4: Busco analysis of translated transcriptomes.

Supplementary Material 5: *L. longissimus* precursors of peptides that were detected by mass spectrometry with highlighted cleavage sites and detected peptide sequences.

Supplementary Material 6: *L. longissimus* mass spectrometry scores including precursor sequences and details about cleavage sites.

Supplementary Material 7: Alignments of MS-peptide 1-5 and RGWa/APGWa precursors. Supplementary Material 8: Analysis of Zieger et al. 2021 deuterostome MIP sequences. Supplementary Material 9: Annotated GPCR trees shown in figure 5, with sequence identifiers on each branch.

Supplementary Material 10: Vasotocin-neurophysin and neurophysin-only trees. Supplementary Material 11-13: Full length sequence database used for GPCR trees shown in figure 5.

Supplementary Material 14-18: Trimmed alignments of GPCR sequences used for trees in figure 5 and vasotocin trees shown in Supplementary Material 10.

Supplementary Material 19-23: Original tree files of GPCR trees shown in figure 5 and vasotocin trees shown in Supplementary Material 10. Root-branches are highlighted in grey.

Supplementary Material 24: Full length reference genes of alignments and information about genomic sequences for intron-exon analyses used in figure 6, figure 7 and supplementary vasotocin trees.

Supplementary Material 25: Collection of pNP query sequence files used for the initial tBLASTn analysis, updated with newly identified sequences. (The dataset is not fully manually annotated and might include sequences derived from spurious clustering.)

### Additional Supplementary Material is provided on external repositories

The mass spectrometry proteomics data have been deposited to the ProteomeXchange Consortium via the PRIDE (Perez-Riverol *et al.*, 2019) partner repository with the dataset identifier PXD023147 (Project name: Neuropeptides in *Lineus longissimus* larvae).

The tBLASTn script, patternsearch script, parsing from phobius analysis script and the general transcriptome assembly pipeline are deposited in the Jekely lab github repository https://github.com/JekelyLab/Thiel_Yanez-Guerra_et_al_2021 (commits: <u>7dd5d28</u>, <u>1b01353</u>, <u>4ed18ea</u>, <u>e6fb954</u>).

Proneuropeptide CLANS file of figure 2 and assembled transcriptomes that were not published elsewhere (Nemertea: *Baseodiscus unicolor, Cerebratulus* spec., *Lineus longissimus, Lineus ruber, Nipponemertes* spec., *Riseriellus occultus*. Brachiopoda: *Novocrania anomala, Terebratalia transversa*. Phoronida: *Phoronopsis harmeri*) are available at zenodo (Project Title: Nemertea, Brachiopoda, Phoronida transcriptomes and proneuropeptide clustermap. DOI: https://doi.org/10.5281/zenodo.4556028).

## Acknowledgements

We thank Silke Wahl from the Proteome Center Tuebingen (Germany) for sample preparation prior to LC MS/MS analysis and Jürgen Berger from the Max Planck Institute for Developmental Biology (Tübingen, Germany) for taking the SEM pictures of the brachiopod, nemertean and phoronid larvae. This research was supported by the FP7-PEOPLE-2012-ITN grant no. 317172 “NEPTUNE” to AH and GJ. The work was also funded by a Leverhulme Trust Research Project Grant RPG-2018-392 to GJ and the NFR FRIPRO Project 288541 “EVOBRAIN” to AH.

