## Supplementary Material 1-25 for "Nemertean, brachiopod and phoronid neuropeptidomics reveals ancestral spiralian signalling systems": Supplementary_Material_05_L_longissimus_MS_precursors.docx

Signal peptide

Cleavage

Amidation + Cleavage

Cysteine

* complete C-terminus

**Detected by MS**

>Allatostatin A1

MGGHARISSLPVILVCVFVNLVLTLCAEINLGSASQDSIQKRDTNSIAALETGRERRSLNSKTNSIKSAKAALGEHVLKRRSLENVEDTSFLDKRAENASPLKRQVDRLLGRDKRDVDKNRFAGGLGKRLDSMMYDVGLGKKL**DPMMFSGGL**GKKL**DPMMFSGGL**GKKL**DPMMFSGGL**GKKL**DPMMFSGGL**GKKL**DPMMFSGGL**GKKL**DPMMFSGGL**GKKL**DPMMFSGGL**GKKL**DPMMFSGGL**GKRVDSMRYSGSLGKRYDQMMFSGGLRKRLDRLMYGGGLDETKKRVDSMKFGGGLGKRLDSMMFGGGLGKRDVGVDKFRFGGGLGKRV**DPMMFSGGL**GKRSKRF**DPMMFSGGL**GKRDKDKR**VDSMRFAGGL**GKRNDDKR

>Allatostatin A2

MTMRPAVFCLLVMVICLARGSTVQQERNDDSGEEATMENSEKSKKDTSEYLSDDSTSDVIDSGAKRGFDRNMYLYGIGKRMEERMPDAGLKR**AMDLFKYRTLGL**GKRGSDYSDDEMKR**GIDHLRYQTLGL**GKR**GIDHVRYQTLGL**GKR**GIDHVRYQTLGL**GKR**GIDHLRYQTLGL**GKRDGDDEDEKR**GIDHLRYQTLGL**GKRDDDKR**GIDHVRYQTLGL**GKRDDDKR**GIDHLRYQTLGL**GKR**GIDHLRYQTLGL**GKRDDMEAEKK**GIDHLRYQTLGL**GKREGIDKK**GIDHVRYQTLGL**GKREGEKRGIDHMKYQTLGLGKREDGEKRGIDHMKYQTLGLGKRDGDDAEKRGIDHMKYQTLGLGKRNDESESSEHTRIER**AVDFIKYRNLGL**GKRDRAYQPYKR**GMDYVLYRNLGL**GKR**AMSDLDLGENDIRPELENA**DLDDESYPMLDMDELVPYIRN*

>Allatostatin C2

MLRSEQYLWIVLFAFLAVCSISQARGIQKRSEGSSFTNQVATKEEHEATGRMLLRLLDAAIEGLREQGDDGNAVTNLVKKR**DYVRDCFFHAVSCY**GRK*

>Allatotropin

MKITLYIFCMAFVVVTIRASPSLRRSKR**SFKDEIDFGHGF**GKRTDHLGNLLKHHTLSDTSDRPLLSNSELAKVIAESRELSEALVAKFVDINGDGYVSKKELFPEGVNL*

>ASWLDF

MRKSQNVLLLFAGLFCLYASVASENKAFEVNGDRISDSVADESKLTEINSGQKRASWLDFKRNGGREYDESDNANQDEAKRASWLDFKRDDAAKKRASWLDFKR**GNTGDLKPWLEN**KRASWLDFKKRASWLDFKRTHGNTDANNDKRASWLDFKRAVDKKASWLDFKRNTDKKASWLDFKRDNDKRASWLDFKRNGASPDDKRASWLDFKR**ALGSWIEAKQT**HKR**GNDNPMDPETIKVI**REMLLKDKSELTPSMRENLLSAFAKDTQK*

>Calcitonin

MQQEFVIKLRYVLLAGLFAVTLA**FEDSRIADAL**RQHKELTAKR**SEIGNVKAVLS**EIDDDLENVQSKICMLSGSTDPACSGYAVKRSDADAADMFSQKDGNRILKDLARKRAALQFLKKLLMDENERNGQVIKR**NCLVNLGGHCATETAA**AIADMYHYLNSAESPGRKKRSASNLLRKALRKN*

>CCAP

MLAYGSVCYILLLTTLISGQESNSQNDFQKNKASGSIFPYGSADNDLDVLKSLRERLDKEKQSRAKR**PFCNSYYGCGNGGY**GKRSEELKISKSEDPSNEQGLLLKALLSNTGNNQFMEGSGKTQKRAFCNGFYGCANGKRSGMENVIQRALKQKTSKMAAKRPFCNAYSGCGNGKKRSVDVENGAEGEKGLLQTILSRLRSVESEGNHQEEPFTF*

>CCWamide

MKLYIVACLFLVCALQFVASYPYGDESKAKSKK**SGDNQENEYGDDQLEVFRDAA**KRAFYNFKCATSSCRPRAMEPTLRCCPGMRCKCGILWNAGQCKCTNYGYGR*

>Corazonin

MEAIVTKSLVLLMVTFTLTQTVHS**QGFHFSQSWRPN**GKRSQMNNNEIMGKVRQWASLGNVENSGKCALTPEGAGAVMKIIQDEVQRSRSFCRSDVAIMSALFNVAGTDIKKD*

>DH44/ELH

MFRHLPFTIGVAIVLLSCLEA**RPYAIQEEYLPLDDEFEADAFRENGDE**KRSMDLSVSGSLAALSDMLMWQRQQQQQQQLNALGKRGSEFNNQETQELLKQLQLLRDSASGMRKRSGSELSVMGSLDALSDIISSQNARHKHDMMQANHQRLLRLGKRDTSANLEKTKNEETKKAD*

>FMRFamide

MYILYLVALLAGQAIATDFNALCSDPKLNSLKNFMVLCDAFRSFSDNLNDVSLDSKRHRATFIRYGRDASHVSQDNALETHLTTPSKRNTEGYMRFGRSVKDPQPAQNVENVDVENTKKEEEKKAIPSTAVDSEQKQETTAEDDKAKRYMRFGRNNYIRFGRESEEADDDEIDDAKRYMRFGKKDDAKRYMRFGKK**NLGPVAEEEGEEQ**KRYMRFGKKDGSDDAEKRYMRFGKRYMRFGKREDDLEEDKRYMRFGKKSDDEKRYMRFGKKDGEEQDEKRYMRFGKKSDEDEKRYMRFGRGDEDMDADKRYMRFGKRYMRFGKRYMRFGKRDEEDANEGGDDEKRYMRFGKRYMRFGKRDTPTDEVRKRNVRSVDESKAEKAAHMRYRR**SEGPDGFERDA**RYMRFGR**NPNGATVDGYIRF**GR**LLDGEDSDETEETPA**KRYMRFGRDYLRFGRLLDGEEESSNDNQNSSPVQKRFIRLGR**LFGGLGVSKEELQ**RRYLRNNRDRAGGYMRFGRSEPAGGYMRFGK*

>FVaFLaFIa

MRSQDILVFCFILGSCLVSTAYADSNDKRARNFLGKR**DSADTKNEDDKSVLNNDDFV**KR**VQEYLDHNDFLEDGLDED**KRGPSFLGKRADDIDEEKRRHSFIGKRRPSFLGKRGPHFLGKRGRYFLGKRGRYFLGKRDGDDEDMDTFDEEKRRLFLGKR**ADLDDTEFGDMEDEM**EKRRLFLGKRVSRNFIGK*

>NKY

MNKYICGCAFLVVLSICYVNSENVDDLHKFLIRRRRSAHNGLRQESELKNLVRELMDKLQNIIDTDRALSSALEEESNEVRHKK**TGFWPSMGPLPVET**R**LSSFGSQIGGATNEQASHKPF**RYGRK*

>Pedal peptide 1

MMFGRGALCLLFLGLACAAPAIDKVEKRSVDRNVLEQLIDSDEKNAAER**ELDTLGGGSIHPETE**RALLDTLGGGAIHPGREVLSSR**ELLESLEEADPEE**RR**ALLDSLGGGAIHPS**RDLDKR**TFDQIGHGGFSSFN**KR**NFDQIGHGGFSAF**NKR**NFDQIGHGGFSAF**NKR**NFDQIGHGGFSAF**NKR**NFDQIGHGGFSAF**NKR**NFDKIGSGSLSSFD**KR**NFDPIGHGGFSSFN**KR**NFDRNGDGSSSGDFA**KR**PFDKIGAGAFSSFS**KK**NFDRIGHGGFSSFN**KK**NFDHIGHGGFSSFN**KK**SFDSIGDGPLSG**FVKRSADKTKEEKKDAE*

>Pedal peptide 2

MSGRVPLSILGLALMACLLSNGVSGEEKRSGSMAGEGQDHKRMLDRIGSGLLKRNGDGEQKR**MLDSISGGLL**KKR**MLDSIAGDLL**KRSGDDQKRMLDRV[……]EKRMLDRVGGRLLKKRWLDHVSGSLLKKRMLDRIGSGLLKKR**LMDPIEEGLL**KKKDAASNDGERRYIDSLSDDLLRRQELNEIIKTRHIDPINANLLRRHLDSIDNDLLRRR**YVDTLGSDLL**RREEAKDEESADEKEE*

>Pedal peptide 3

MRVLPNIVLLSLQLLLISRLTFQ**EKIQTDSGVDI**KRVER**SVDDGENSFGSED**KRVFGSITKPVGKHGSARRPPATREEHVRVSDIKREDEPDYYKLLDELMDRRIESKRAAYDANAGDEGDGGSPDDEGNNMLYLALKQGEHNDNGDDDGVLHSYDKR**PFGSFTRKLS**KR**PFGSLTGGFH**KR**PFGSFTRKLQ**KRPFGSMTGQMKR**PFGSIISGLS**KR**PFGGFTSRM**KK**PFGSLSKVLS**KKPFGSFSRPRSFNKREDEDVEKRPFGSFSRPRWNTKRDDDNDVEKRPFGSFSRPRWNTKRDDDNEVEKRPFGSFSRPRWNTKRDDDNEVDKR**PFGSFSLPR**SYQKREGDDMAKR**PFGSFTRPNT**RSF**PTEDDTDDIIE**KRPFGSFTRWRSRTRGKR**DGADEVEHDTPVYY**

>GNXQN/Prohormone 2

MMVFRALLALTLVLCLTQALPLKKANDLSKEIADVAKDNTQKRLVKR**SEETVIVGNHQNAPRPALEL**KNRDENGKIEVKENEVDKTEIAKSDVADDTPPDDEVMEKVVMDLAKVAEAVPETDNNSDLEEKVKVDEDEKVLDEVPNNDEVAKEKEVIEEDAVFQPQEIFNLGAQEKEINDVNSSELPDQESEVTESSEEVPPEELMSGNMEMAQEQPAADEYQYDEYNALYSDPYQWLDRKKRNVNRQKREAALQGSEAGMSMVANEDDTKSRSKRDTEYPLSLNDLYNLYYKYKEMEDEKQQEEELEPVYELPEYEENYAANPEYDNYGVEEELAPPSEEWEPAAVNDDIQQELELEAMMEPAMEEEAPYYLADDETPEEELEKEMEEELEEQMEAQQLPAYDVPSEPYYYPMEYNPIAPQPDKR**NQEMLSMLPGI**KRADDFYPSYTEDREPWQALIPPAAEKRGVMEEYARLYRLARALKRSREDSVEERWESLLGDLDEKKK*

>Prohormone 3

MMKAVLCFGLLAILIGCVRGW**GRILSGENSIGYNS**WIPARRSWRCGGNRDLCWQNSQCCKGYYCATVDDGKSGFCRAEDQEQIPVCETDSDCPGWTKCSTVAQVGAVKLRMCKEPTDTAADTGTGKKAESGQPGDVCEDNSDCSIADGLCCQYVQVFRRKPKKMCHQISGLNKCIKSTSFGNNIIKK*

>PSLamide

MGVYDKRWLVSWHNMVLAALYCLFLATTAVTSSEVSNGENWDTVLSDSEMANSPAESQENSIEDSLNLTNNDYDMDIENDSAEFPIGGANDESSAAKRFSLTAPSLGKRFSLTAPSLGKRFSLTAPSLGKRFSLVTPSLGKRFSLTAPSLGKRFNLVSPSLGKRLRHALVSPAFGKR**DEEEYADRLDSDDEPRDES**KRYSLVTPSLGKKMSGIVAAALGKRDAEKRFSLTAPSLGKRFSLTAPSLGKRFSLTAPSLGKRFSLTAPSLGKRFSLT

>RGWamide

MKLYIAITLVCLLVFEIFASAAAAETADQDMDKRAPGWGKRGWGKR**DDDSEGALDDYL**KRGWGKRGWGKRGWGKRGWGKRGWGKRGWGKRDIDGAPCEDLNQGVIYYIYKAVEAEAQRINACASTDN*

>SIFamide/FF peptide

MNCYVKLSIILCFCSLLVCG**QASQGIDKLLKSPNLL**FGRKR**SPDPAGSNLFF**GRRSFEPQQYGRQFCANVIETCRYWYEQNNDVSATEQS*

>MS peptide 1

MMRQCFGPFLIGLIGILICFTQDARG**SPLPDNEEMVNEILN**KRNAWWSKKSDPALADLSGERMDGENSEQPCLSQAHRIDCYINRCVPDFVECGRLARSRSGFGACKQQHHECALECYPDTNKGPALSLFSGLQG*

>MS peptide 2

SYLVACIVVLCACLVLGKPAPSGTKVNSQDISKSINRLIEDLRTHKAAEDHDQKELVKIVLAFAGKLQELQEGIQHIDEVEGQAAKATAKR**EGSEGGDFLERLD**KR**GQYYDAELDDGFDMF**KKRNQKLNRARNGWLEQ*

>MS peptide 3

MTRIYALLAVALLVTQVLSAPVAKQDVGERKAKNKEAANLEEELHKQAGEGKKDTPETSEPSPGHKPRGALGQLMQAAAEEEESQEEKAATTKTEEDAKLQNVIYSIYNNPEVLKALLRAGETKESNEEVAPPMQNGIKR**DDEEEELRSNII**KKDGDDLYQPASDPNMDMLLDRYEDLPDDSALYEDDRQYNNVADPDYPDDRLMAEEVHALEGMEPNMNGIEVEDENEKDEDDSEDHDDNEEESTTKSASKQ*

>MS peptide 4

MDGRLLFLVIALSACGSLLACKLPREPTQGEYMKYIVCLQKATGFTSDGGRWGR**SVENVLDRSQPL**RKLLRSKKTECVLKKNPQPQDYVDYVMCLHSNGHTTEGGRWGRSVDEGLTRDGLKSKSKREECVLKSNPQPQDYVNYVMCLQGTGFTT[……]TTDGGRWGRSIDREFTRNGLKSKSKRDQCVLKSNPQPQDYVNYVMCMQGSGFTSDGGRWGRSTDRARHYSIMEKVFRNVPDFGRWFQLRRESAHEQAEIPARQGGRDDDQETVFSR**QANGRVQDLANAGVTIIEHPV**KREIRLRDITAKKSS*

>MS peptide 5

MDTCMKVLFCLYAISMLMMATSRVTNALSSDFGGKPQRR**EATRGNLMPPGPWIYGTGHLN**KRLFKGAFDISNKIEDTAGKEHGIQMMDKFMVFLRLQESGALDKCLASFSKRR*

>SWD peptide, C-terminal fragment

ERSARPTQKR**QYEGIFITDGSQ**KRSWDALGIPDKRSWDALGIPDRRSWDALGIPDRRSMGLNEKDVKALASFIASRNNRRQ*
