## Supplementary Material 1-25 for "Nemertean, brachiopod and phoronid neuropeptidomics reveals ancestral spiralian signalling systems": Supplementary_Material_07_Alignments.pdf

### WWS peptide (MS peptide 1)

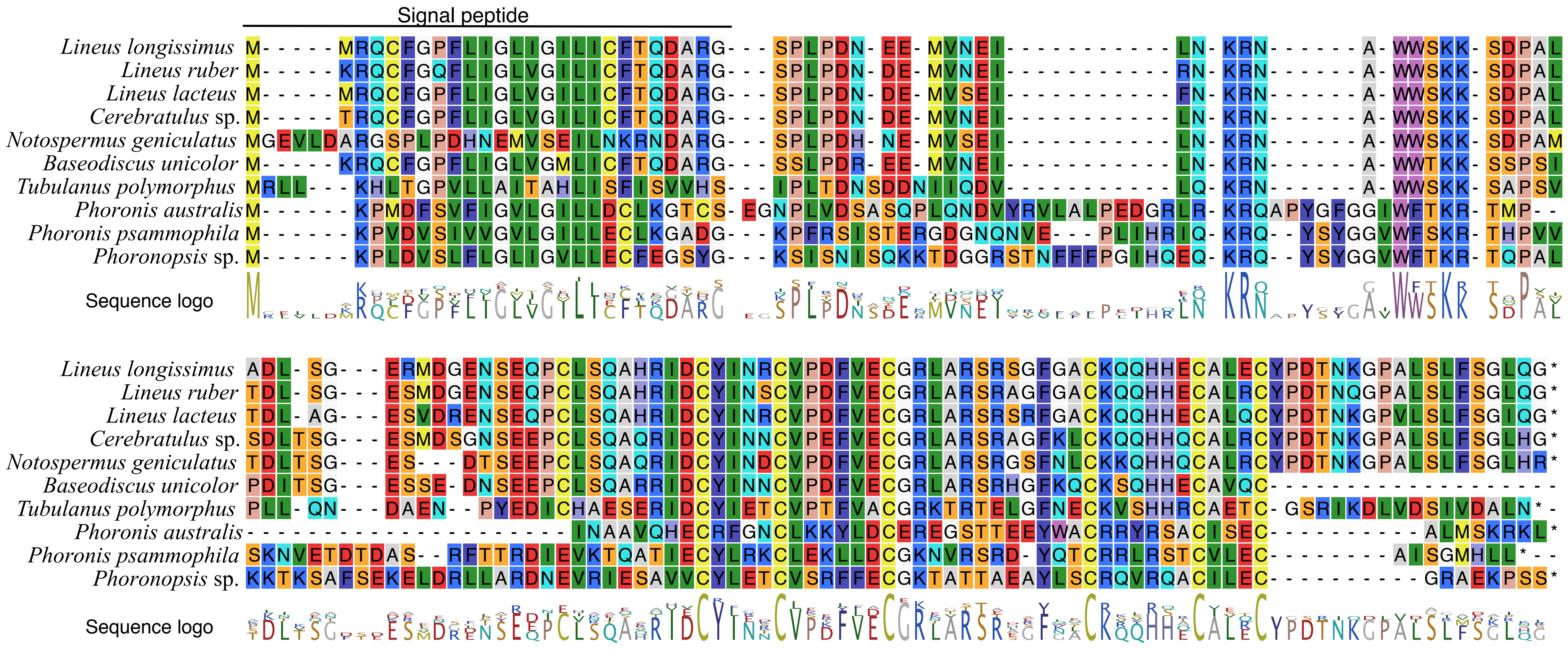

### DMF peptide (MS peptide 2)

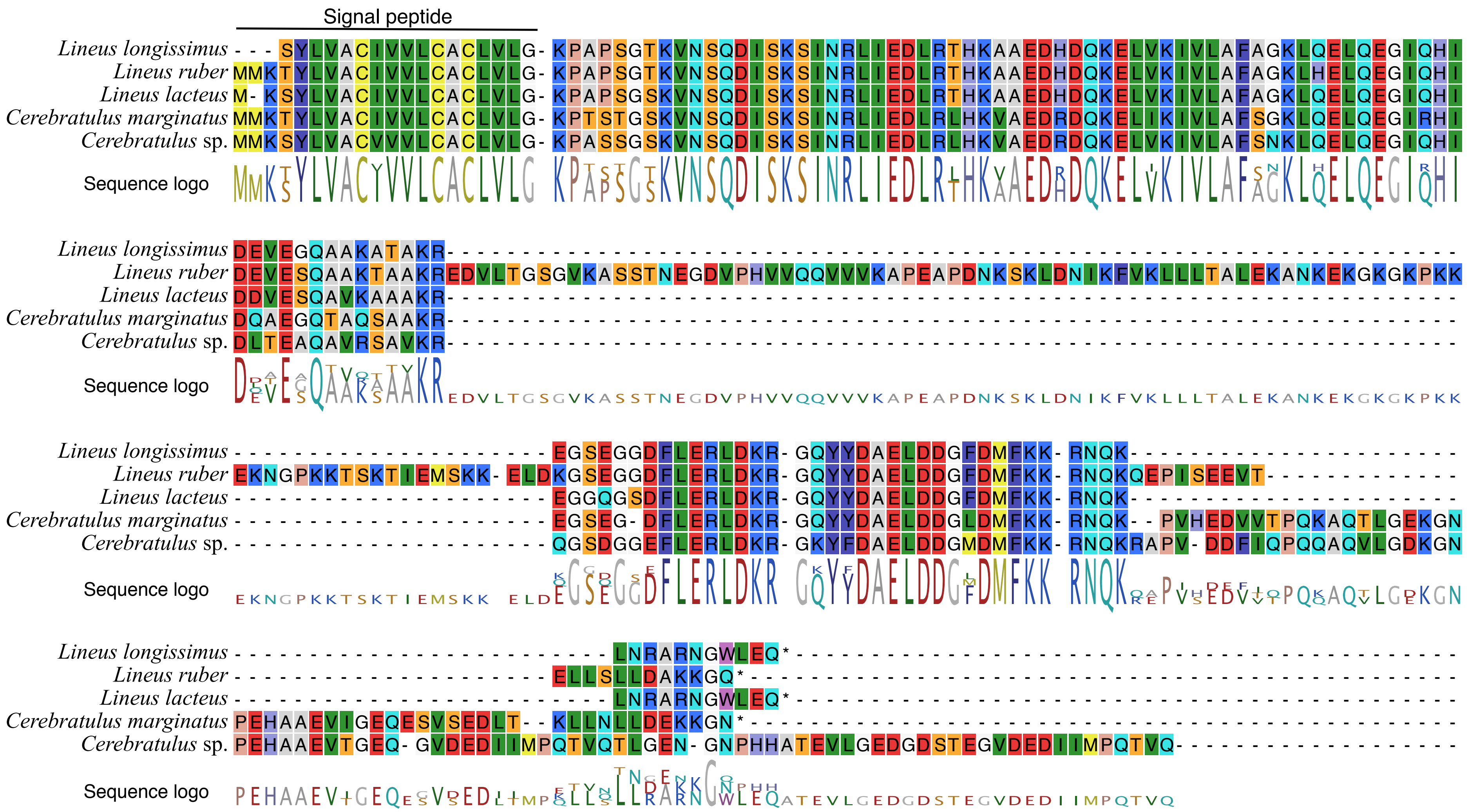

### AGEamide (MS peptide 3)

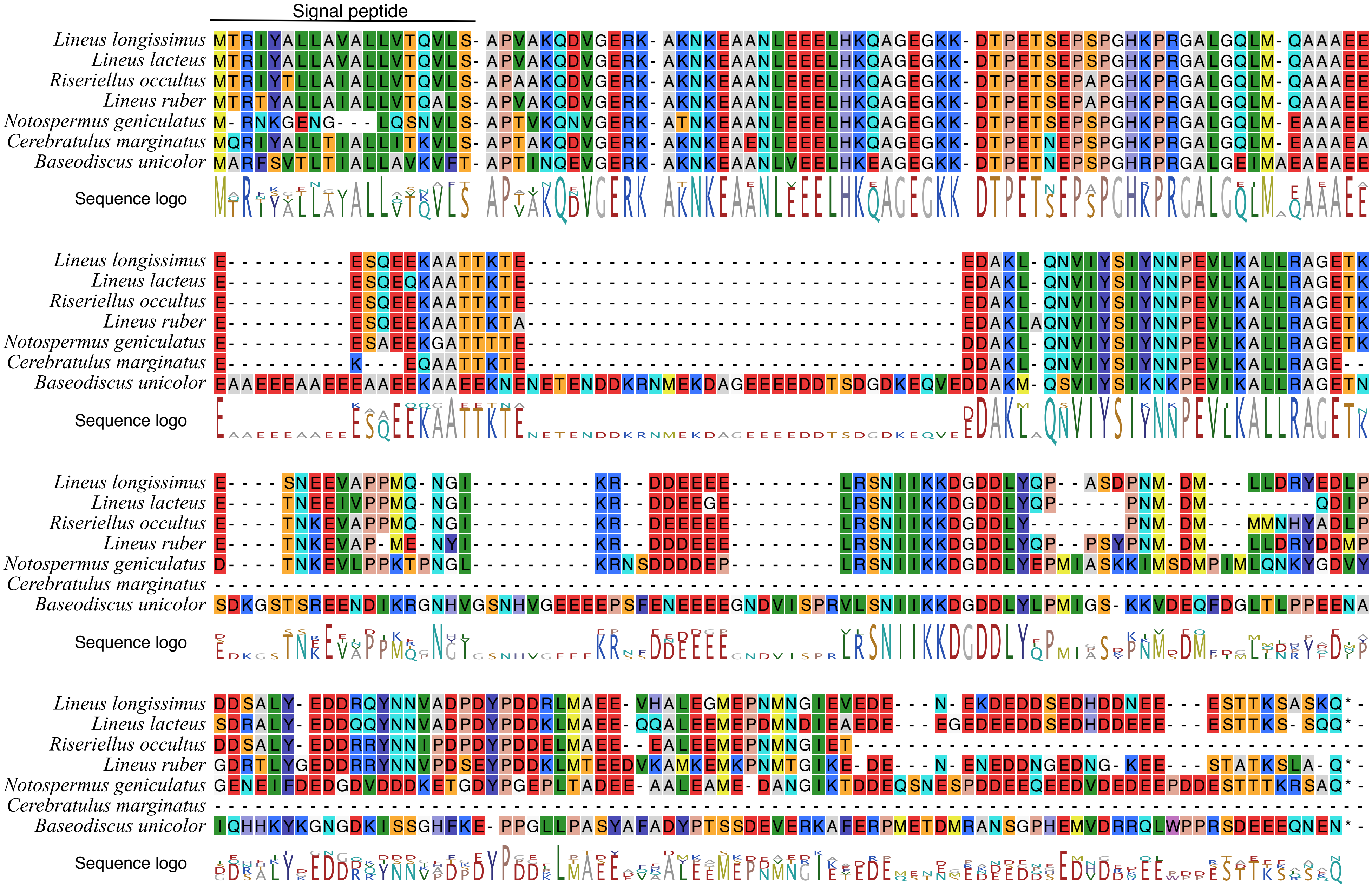

### GGRWamide (MS peptide 4)

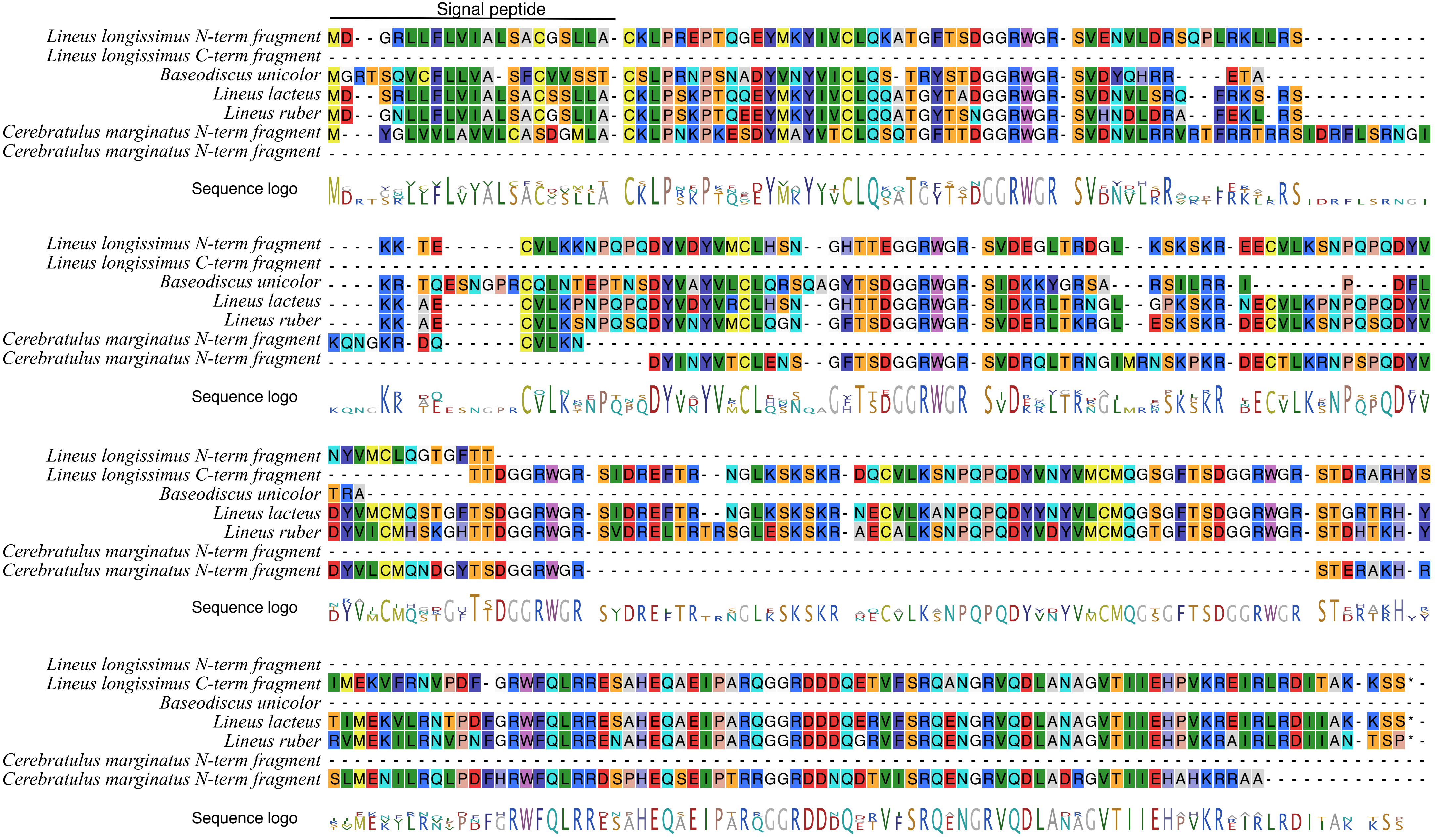

### GxGH peptide (MS peptide 5)

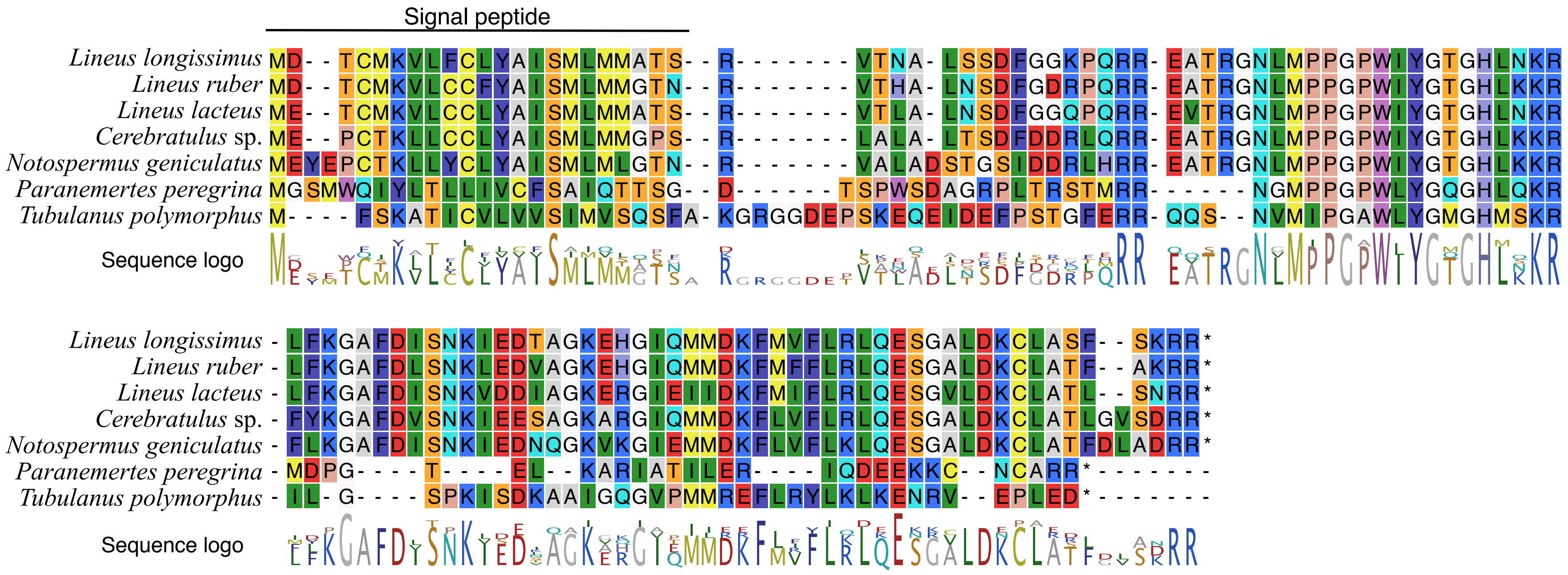

### RGWamide/APGWamide

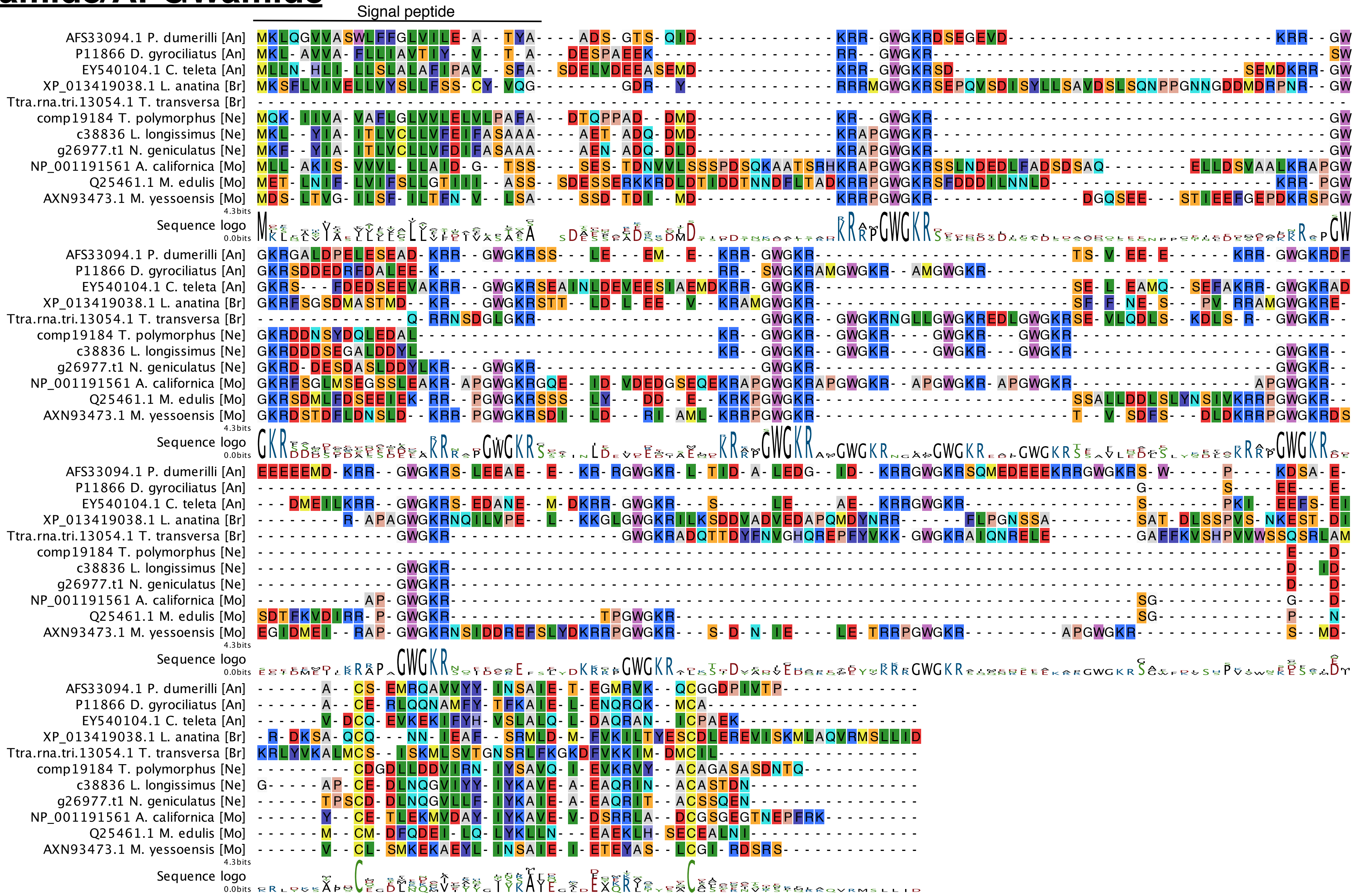
