## Supplementary Material 1-25 for "Nemertean, brachiopod and phoronid neuropeptidomics reveals ancestral spiralian signalling systems": Supplementary_Material_08_ZiegerEtAl2021_MIP_analysis.docx

Comments on the alleged deuterostome MIP precursors reported in

Zieger E, Robert N, Calcino A, Wanninger A **(2021)**: Ancestral Role of Ecdysis-Related Neuropeptides in Animal Life Cycle Transitions. *Current Biology 31 (1)*: R30-R32.

We were surprised to read in this paper that the authors identified two ambulacrarian myoninhibitory peptide (MIP) precursor sequences. MIPs or MIP receptors could not be identified in previous large-scale bioinformatic studies (Mirabeau and Joly 2013; Jékely 2013; Williams 2020).

Proctolin and MIP receptors were previously found to be many-to-many orthologs to deuterostome GPR139 and GPR142 receptors (Mirabeau and Joly 2013). Another study of GPCR phylogeny with an increased taxon sampling failed to identify a one-to-one MIP receptor ortholog in ambulacrarians (Thiel et al. 2018). Given the strong coevolutionary pattern of neuropeptide and receptor systems, this absence suggests that deuterostomes lack MIP signalling and their receptors are members of an extended protostome-specific signalling system.

We therefore read the new findings by Ziegler et al. with scepticism and had a closer look at the reported MIP-related proneuropeptides. The paper reports two sequences as ambulacrarian MIPs, one from the hemichordate *Saccoglossus kowalevskii* and another one from the sea cucumber *Apostichopus japonicus.*

Below we show that these sequences are misidentified as MIPs and very likely do not even represent proneuropeptide sequences.

MIPs are characterised by two W residues, one of which is at the C-terminus of the peptide and is amidated and another one close or at the N-terminus of the peptide with usually 5-8 amino acid residues in between. These two W residues are important for peptide activity and stabilize a beta-turn conformation (Kim et al. 2010).

Sequence 1 (Saccoglossus kowalevskii):

>SACKO_MIP-like_Sakowv30027671m

MPPVMSALENDTLLQTSDRNLDYNGTEKHEWGKQTEEAGVAQWRKQTEEARVAQWGKQTEEAGVAQWGKQTEEARVAQWGKQTEEARVAQWGKQTEEARVAQWGKQTEEAGVAQWGKQTEEAGVAQWGKQTEEAGVAQWGKQTEEAGVAQWGKQTEEAGVAQWGKQTEEARVAQWGKQTEEAGVAQWGKQTEEARVAQWGKQTEEAGVAQWGKQTEEAGVAQWGKQTEEAGVAQWGKQTEEAGVAQWGKQTEEARVAQWGKQTEEARVAQWGKQTEEAGVAQWGKQTEEAGVAQWGKQTEEAGVAQWGKQTEEAGVAQWGKQTEEAGVAQWGKQTEEARVAQWGKQTEEAGVAQWGKQTEEARVAQWGKQTEEARVAQWGKQTEEARVAQWGKQTEEARVAQWGKQTEEARVAQLKHAHKAFLKLASKIKWKIVMLF

Full-length proneuropeptide sequences are characterised by an N-terminal signal sequence. We failed to identify a signal peptide in this *Saccoglossus kowalevskii* sequence (with SignalP 4.1 or 5.0).

Proneuropeptides also have characteristic cleavage sites, most often dibasic sites (e.g. KR), and the individual peptide sequences are often interspersed with acidic intersequences (also in MIPs). There is only one dibasic site in the whole sequence and no consistent basic amino acid N-terminal of the “WGK” motifs to support a general alternative monobasic cleavage site at the “K” (only in some cases an “R”). The whole sequence only possesses one cleavage site according to “NeuroPred”. Even if these sequences were cleaved at these monobasic sites, the resulting peptides would only possess a single “W” residue, whereas MIPs/allatostatin-B/prothoracicostatic peptides are characterised by two tryptophan residues. In addition, the sequence is highly repetitive and contains 30 identical VAQWGKQTEEA stretches, a level of repetition and conservation that is highly unusual for a proneuropeptide.

Sequence 2 (Apostichopus japonicus):

>APOJA_MIP-like_MRZV01000188.1_790888:792222(+)

MLELASALLSLSQKNKIQTAMEMRSDSLPHCTWGTRILDSLPHCTWGTRRLDSLPHCTWGTRILDSLPHCTWGTRMLDSLPHCIWGTRRLDSLPHCIWGTRILDSLPRCIWGTRMLDSLPHCIWGTRILDSLPHCIWGTRIIDSLPHCIWGTRILDSLPHCIWGTRILDSLPHCTWGTRMLDSLPHCTWGTRIIDSLPRCTWGTRRLDSLPRCTWGTRRLDSLPRCTWGTRILDSLPRCTWGTRRLDSLPRCTWGTRILDSLPRCTWGTRRLDSLPHCICGTRIIDSLPHCTWGTRIIDSLPHCTWGTRMLDSLPHVHGNTNNRFLATLYMGNTKTRFLATLYSGNTKTRFLATLYMGNTNTRFLATLYMGNTNTRFLATLYMGNTKTRFHKNIQSLLVTSKKLALMWNSMYCMIIAVLFDLWC

There is no indication of an N-terminal signal peptide (tested with SignalP versions 4.1 or 5.0). There are only a few cleavage sites according to “NeuroPred”. If there is alternative monobasic cleavage at “R” it would be supported by the “histidine” 6 residues N-terminal of the cleaved “R”, which is very unusual. Even if there is monobasic cleavage then the resulting peptides would only possess a single W residue whereas MIP peptides have two tryptophan residues. This W residue would also not be at the peptide C-terminus and would not be amidated.

A Pfam motif search (<http://pfam.xfam.org/search/sequence>) of this protein suggests a potential homology with Cyclin M transmembrane N-terminal domain-containing proteins. The analysis of this sequence with Phobius reveals a transmembrane domain in positions 405 to 423.

In conclusion, neither of the two sequences are MIP precursors.

Furthermore, several other sequences in the MIP alignment shown in the paper, are actually not MIP but APGWamide/RGWamide or YWamide precursors. The mis-identified MIP-related precursors contains mis-identified precursors include the RGWamides SCHME_MIP-like_SMEST015527001.1, CRAGI_MIP_EKC38991, HALDIHA_MIP_HDSC01375CG00010, PERAI_MIP1_GDAF01004548.1 and a YWamide peptide APLCA_MIP2_XP_005107870.2_XM_005107813.2.

The clustering of the RGWamide sequences, the YWamide and the two non-MIP ambulacrarian sequences with the bona fide MIPs can simply be explained by spurious pattern matches of these repetitive sequences during the Clans analysis. The Clans analysis was also carried out improperly, with only a very small number of sequences containing a pre-selected set and therefore precluded an unbiased global analysis of sequence similarities (Frickey and Lupas 2004).

Williams, Elizabeth A. 2020. “Function and Distribution of the Wamide Neuropeptide Superfamily in Metazoans.” *Frontiers in Endocrinology* 11 (May): 344.
