## Supplementary Material 1-25 for "Nemertean, brachiopod and phoronid neuropeptidomics reveals ancestral spiralian signalling systems": Supplementary_Material_09_Annotated_GPCR_trees.pdf

a)

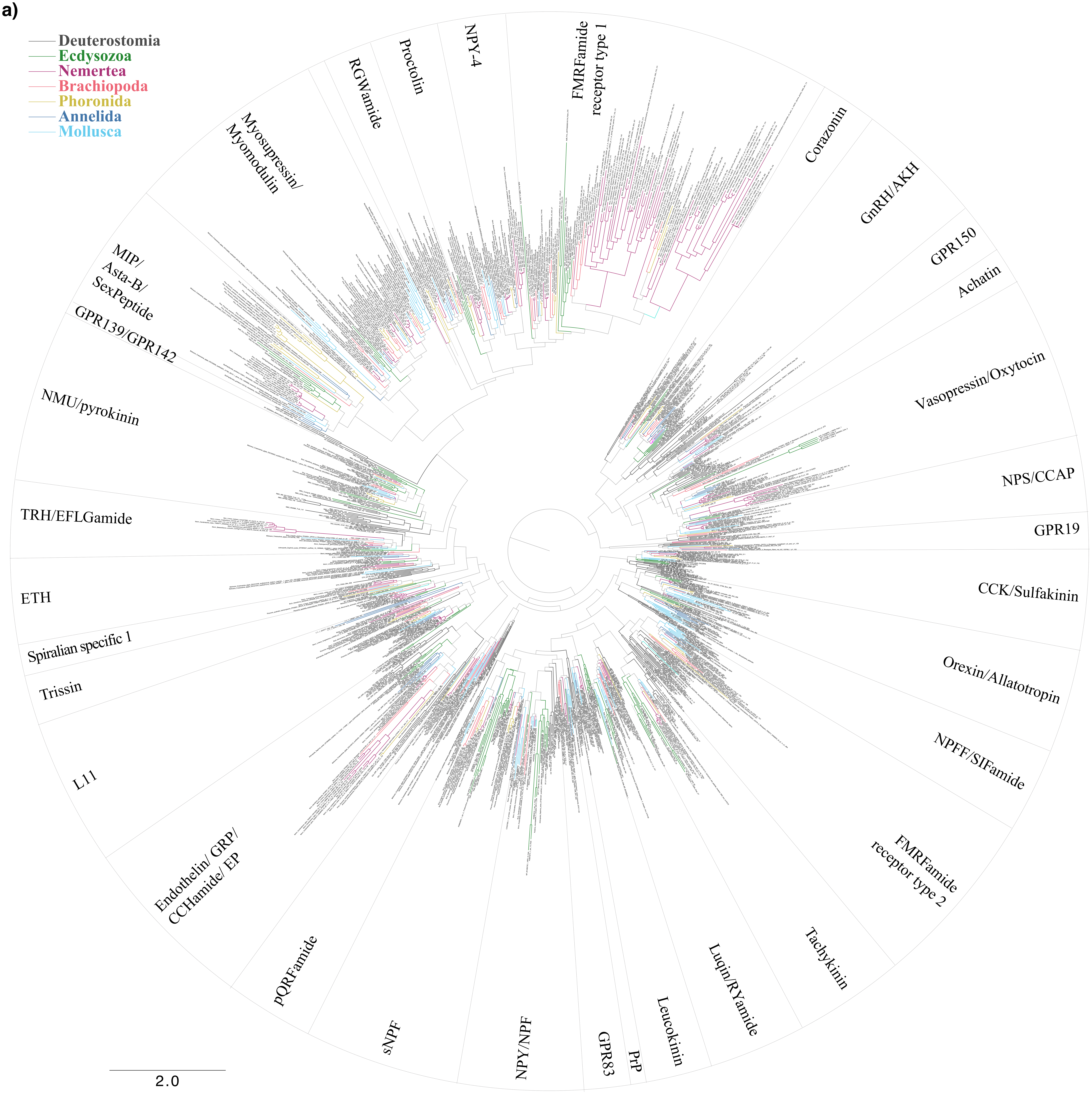
Deuterostomia
Ecdysozoa
Nemertea
Brachiopoda
Phoronida
Annelida
Mollusca
FMRFamide receptor type 1
NPY-4
Proctolin
RGWamide
Myosupressin/ Myomodulin
MIP/ Asta-B/ SexPeptide
GPR139/GPR142
NMU/pyrokinin
TRH/EFLGamide
ETH
Spiralian specific 1
Trissin
L11
Endothelin/ GRP/ CCHamide/ EP
pQRFamide
sNPF
NPY/NPF
GPR83
PrP
Leucokinin
Luqin/RYamide
Tachykinin
FMRFamide receptor type 2
NPFF/SIFamide
Orexin/Allatotropin
CCK/Sulfakinin
GPR19
NPS/CCAP
Vasopressin/Oxytocin
Achatin
GPR150
GnRH/AKH
Corazonin
2.0

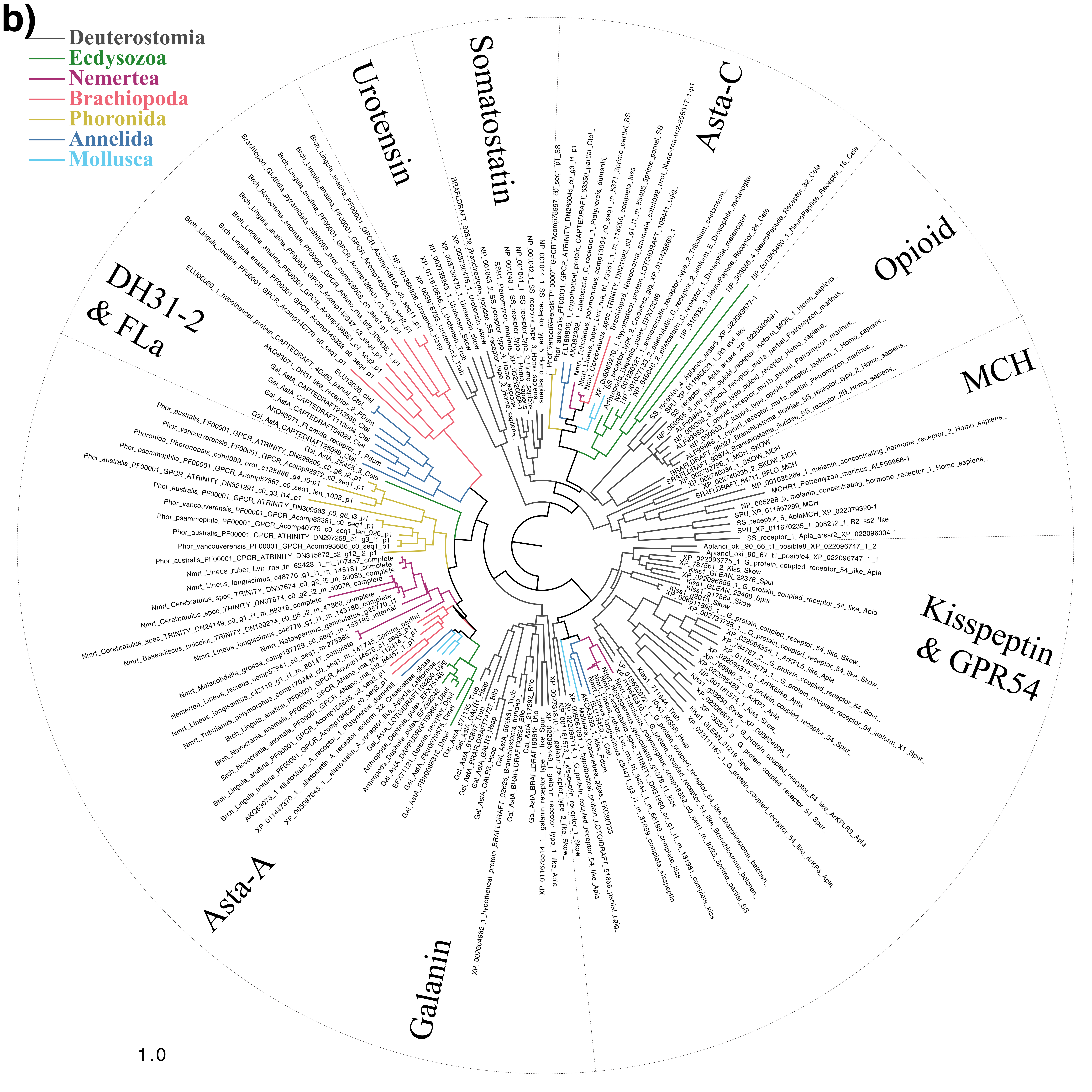
b)
Deuterostomia
Ecdysozoa
Nemertea
Brachiopoda
Phoronida
Annelida
Mollusca
Urotensin
Somatostatin
Asta-C
Opioid
MCH
Kisspeptin & GPR54
Galanin
Asta-A
DH31-2 & FLa
1.0

c)

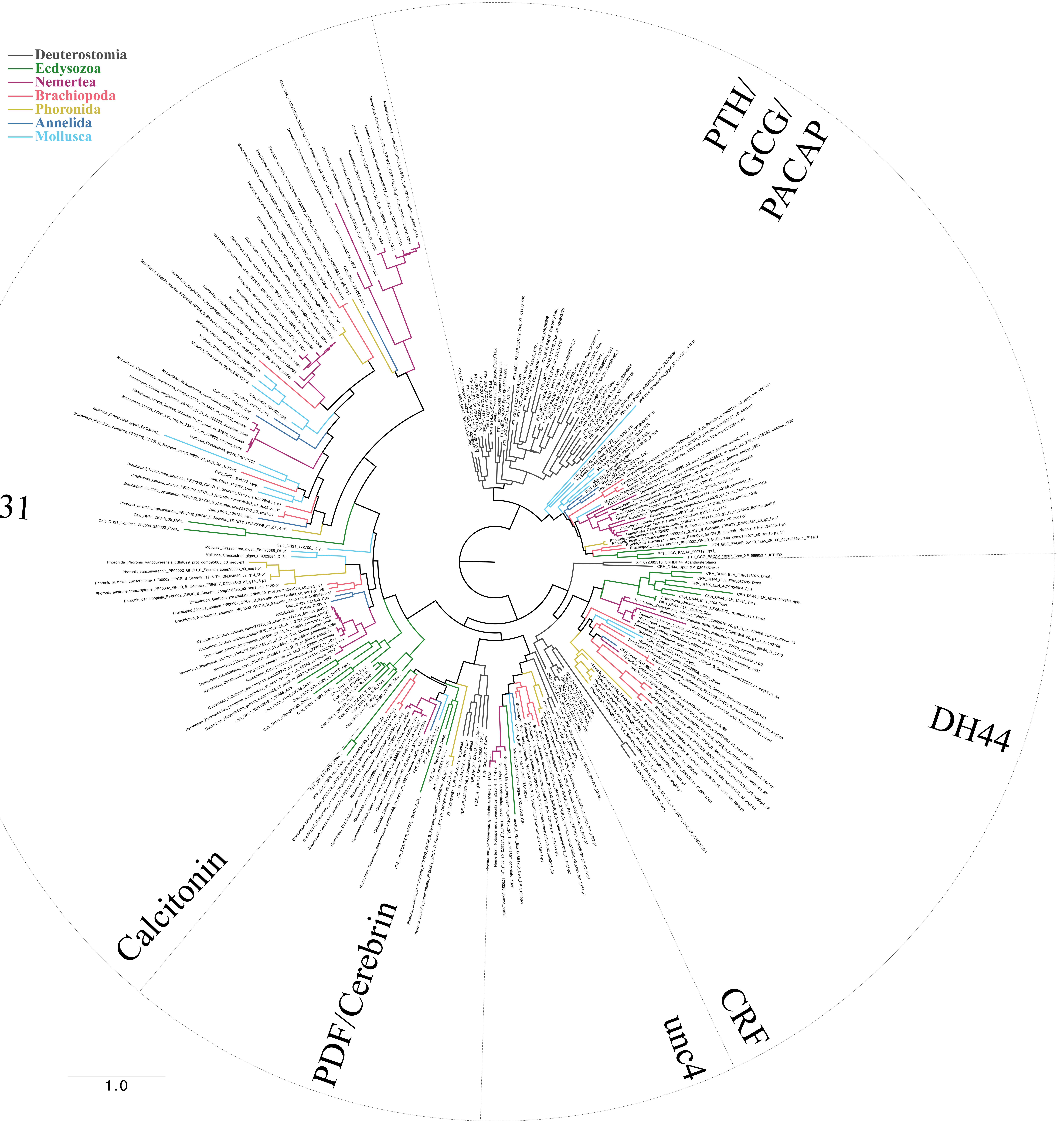
Deuterostomia
Ecdysozoa
Nemertea
Brachiopoda
Phoronida
Annelida
Mollusca
PTH/GCG/PACAP
DH44
CRF
unc4
PDF/Cerebrin
Calcitonin
DH31
1.0

**Supplementary Figures: Maximum-likelihood analysis of rhodopsin and secretin-type neuropeptide GPCRs. a)** Rhodopsin *beta* GPCRs. **b)** Rhodopsin *gamma* GPCRs. **c)** Secretin GPCRs. Terminal branches are colour-coded according to taxon as shown on the upper left. Scale bars on the lower left of each tree indicate the inferred amino acid substitutions per site. Dashed lines demarcate orthologous receptor types. AKH, adipokinetic hormone; Asta, allatostatin; CCAP, crustacean cardioacceleratory peptide; CCK, cholecystokinin; CRF, corticotropin releasing factor; DH, diuretic hormone; ETH, ecdysis triggering hormone; EP, excitatory-peptide; GCG, glucagon, GHS, growth hormone secretagogue; GnRH, gonadotropin releasing hormone; GRP, gastrin releasing peptide; NMU, neuromedin-U; NPS, neuropeptide-S; NPY/NPF, neuropeptide Y, neuropeptide F; MCH, melanin concentrating hormone; MIP, myoinhibitory peptide; PACAP, pituitary adenylate cyclase-activating polypeptide; PBAN, pheromone biosynthesis activating neuropeptide; PDF, pigment dispersing factor; pQRFPa, pyroglutamylated RFamide peptide; PTH, parathyroid hormone; PRP, prolactin releasing peptide; SH, Shimodaira–Hasegawa approximate-likelihood ratio test; TRH, thyrotropin releasing hormone; VIP, vasoactive intestinal peptide.
