## Supplementary Material 1-25 for "Nemertean, brachiopod and phoronid neuropeptidomics reveals ancestral spiralian signalling systems": Supplementary_Material_24_Reference_genes_for_alignments.docx

Signal peptide

Cleavage site

Amidation site

Cysteine

**Reference Precursor sequences:**

**Vasotocin**

>ELT90472.1 hypothetical protein CAPTEDRAFT_173251 [Capitella teleta]

MRHSSDAVNSVVLCGRLLLLFACISCCIETTSGCFIRNCPIGGKRSSVPSRISAQKECMACGPNGLGQCVGPNTCCGQDIGCFMGTQEAKMCGEENDSPIPCRVDGAACGRNDGGRCVAEQICCNEDKCSHDSSCQSKAKRQHDNLSQDLLRYMHQLMTLKSLGGRR

>ABR68852.1 vasotocin-neurophysin, Platynereis dumerilii

MQFSRPTFTLQYGSAVLLTLVVCCSACFVRNCPPGGKRSMDLPQIHSTRQCMRCGPQGLGQCFGPNICCGPSIGCYINTLESEECSKENDVRTPCDITAEICGVDGQGRCGADGVCCTDEKCTLDSSCDKLDMDKPRFSPDILRMLKKIFDRRSYPGRRK

>BAA36458.1 Annetocin precursor [Eisenia fetida]

MACTKKSANMKLRKSLTVTAFLLFVNLSLSSACFVRNCPTGGKRSVLLSPLQPARQCMPCGATVGGRSVLGVCVSENTCCVAHLGCFVNTEESKVCALENHLSTPCRLEGPPCGSDGQDVCAVEGICCAGQNCRYDAQC

>XP_009016584.1 hypothetical protein HELRODRAFT_77987 [Helobdella robusta]

MISNNFFFITSIFFVLLTLEPNNACFIRNCPIGGKKRSYEDDGTNFRQCPPCGTSGPENNLQGICVSENL

CCIDNIGCFRNRSLTADCKVENMIPSPCVLIGKKCGKYDGVCAAKGVCCNNNG

>XP_011434624.1 terepressin/terephysin, Crassostrea gigas

MQKMECCKILHLSLPQVSLLLLMFTTLSSCCFIRNCPPGGKRSMGIVSQPTKECMSCGPGLLGQCVGPDICCGPFGCQMGTSESNICGKENESTTACAISGPPCGSRNQGNCVADGICCDTGACSFNTKCKLNSEPRDVQILSLLKTLGLIETKAYNNAGMS

>AQS80498.1 Conopressin precursor [Charonia tritonis]

MQGYGMKFSVVPVVLFMLLASSYGCFIRNCPRGGKRAFDGGKPCMPCGPDGAGQCVGPAVCCGKSFGCLVGTREARECEKENESSTACSVQGRQCGRDNSGRCVAKGICCVADACSFNERCAEIERNGRDELLGLIRRLLVTHQYE

>XP_021350846.1 terepressin/terephysin-like [Mizuhopecten yessoensis]

MEWRLYVSTRLCLTCVLLCVLVLDCQGCFIRNCPPGGKRSMGMMTRATHQCASCGPGMRGQCVGPNTCCGDFGCLMGTEESRVCMKEDDSTEACAVRGSSCGSMGQGNCVSEGLCCDAYACSYNNKCKVNSGKEDREILSLLNRLLHTNDYTD

>NP_001191416.1 Lys-conopressin preprohormone precursor [Aplysia californica]

MSPLSVRTFVLVAGLAVISFSVVADACFIRNCPKGGKRSMDMQLLGQRQCMACGPEGIGQCVGPNICCSPHFGCHIGTPETEICQKENQSTSPCSVRGETCGYRDSGNCVANGICCDSESCAANDRCRLRKETSRIGFDDTQSSRAEVLKLIQKLLRAKEED

>XP_011065328.1 PREDICTED: neurophysin 1-like [Acromyrmex echinatior]

MLKELIVFASLIFLSYACLITNCPRGGKRSDIASLKTVIRECPSCGPNHLGQCFGPYICCGPSIGCFIGTPETFRCRKESLYTRPCIAGYAMCRGKTARCASDGICCSQASCHMDTSCKISDVGNDRNLDDNVNVILPGNEVSNEILQ

>XP_011147500.1 neurophysin 1 [Harpegnathos saltator]

MLRELVVFASLIFLSYACLITNCPRGGKRGDIIPSLGTVTRECPPCGPNHLGQCFGPHICCGPTIGCFIGTPETYRCRKESPYARPCIAGYAMCRGNTARCATNGICCSQDSCHMDTSCRISDVVSNDRKMDADLSAILSSNEASHEIIQ

>XP_001606547.1 neurophysin 1 [Nasonia vitripennis]

MSKVIIVLTTLVALSYGCLITNCPRGGKRGDPTFLLENIARECPACGREEQGRCFGPHICCSPSMGCLIGTPETLRCRKESLYSRPCVAGFAMCQGNSGRCAANGICCSQESCFIDSACKLVDETGNDRKIGAEFGAFLLENAGTNEHIL

>XP_001606547.1 neurophysin 1 [Nasonia vitripennis]

MSKVIIVLTTLVALSYGCLITNCPRGGKRGDPTFLLENIARECPACGREEQGRCFGPHICCSPSMGCLIGTPETLRCRKESLYSRPCVAGFAMCQGNSGRCAANGICCSQESCFIDSACKLVDETGNDRKIGAEFGAFLLENAGTNEHIL

**GnRH**

>AHB62361.1 GnRH1 precursor, exAKH1, Platynereis dumerilii

MRAWLVLLVACIVIAYQAHTSDAQFSFSLPGKWGNGKRAALGWGKRGECGDFDPDAIFNVYRAIQAEALRINECMQQKLEEESKKH

>AHB62375.1 GnRH2 precursor, ex AKH2,Platynereis dumerilii

MMRLWVVYLTVCVMMLFAFFPHVCHSQLTQSLGWGSAGSSGKRSVKSPIYYGDDDDYDYDVRQVRKNRLTLSEALCSQDEKDAISKIKKLIQREVVRQRYCNRKK

>NP_001104825.1 AKH 1 precursor, Bombyx mori

MYKFTILFLVLACFIMAEAQLTFTSSWGGKRAAIAGTVSCRNDESLASIYKLIQNEAEKLLLCQKP

>NP_001124365.1 AKH 2 precursor, Bombyx mori

MGRALVLVLILSAALLVCEAQLTFTPGWGQGKRSEATDYRNDGCSSEDSVYTIYKLIKNEAEKFLACRS

>NP_001127713.1 ACP precursor, Bombyx mori

MMKVKIFRPYFSSFFVWAACALAAVAAQITFSRDWSGGKRSVAEAPVDCRQFTRFCRHFVVSMQPHHSICSSGPRTQLGSR

**RGWamide/APGWamide**

>AFS33094.1 RGWamide Platynereis dumerilli [Annelida]

MKLQGVVASWLFFGLVILEATYAADSGTSQIDKRRGWGKRDSEGEVDKRRGWGKRGALDPELESEADKRRGWGKRSSLEEMEKRRGWGKRTSVEEEKRRGWGKRDFEEEEEMDKRRGWGKRSLEEAEEKRRGWGKRLTIDALEDGIDKRRGWGKRSQMEDEEEKRRGWGKRSWPKDSAEACSEMRQAVVYYINSAIETEGMRVKQCGGDPIVTP

>P11866 RGWamide Dimorphilus gyrociliatus [Annelida]

MKLAVVAFLLIAVTIYVTADESPAEEKRRGWGKRSWGKRSDDEDRFDALEEKRRSWGKRAMGWGKRAMGWGKRGSEEEACERLQQNAMFYTFKAIELENQRQKMCA

>EY540104.1 RGWamide Capitella teleta [Annelida]

MLLNHLILLSLALAFIPAVSFASDELVDEEASEMDKRRGWGKRSDSEMDKRRGWGKRSFDEDSEEVAKRRGWGKRSEAINLDEVEESIAEMDKRRGWGKRSELEAMQSEFAKRRGWGKRADDMEILKRRGWGKRSEDANEMDKRRGWGKRSLEAEKRRGWGKRSPKIEEFSEIVDCQEVKEKIFYHVSLALQLDAQRANICPAEK

>NP 001191561 APGWamide Aplysia californica [Mollusca]

MLLAKISVVVLLLAIDGTSSSESTDNVVLSSSPDSQKAATSRHKRAPGWGKRSSLNDEDLFADSDSAQELLDSVAALKRAPGWGKRFSGLMSEGSSLEAKRAPGWGKRGQEIDVDEDGSEQEKRAPGWGKRAPGWGKRAPGWGKRAPGWGKRAPGWGKRAPGWGKRSGGDYCETLEKMVDAYIYKAVEVDSRRLADCGSGEGTNEPFRK

>Q25461.1 RPGWamide Mytilus edulis [Mollusca]

METLNIFLVIFSLLGTIIIASSSDESSERKKRDLDTIDDTNNDFLTADKRRPGWGKRSFDDDILNNLDKRRPGWGKRSDMLFDSEEIEKRRPGWGKRSSSLYDDEKRKPGWGKRSSALLDDLSLYNSIVKRRPGWGKRSDTFKVDIRRPGWGKRTPGWGKRSGPNMCMDFQDEILQLYKLLNEAEKLHSECEALNI

>AXN93473.1 APGWamide Mizuhopecten yessoensis [Mollusca]

MDSLTVGILSFILTFNVLSASSDTDIMDKRRPGWGKRDGQSEESTIEEFGEPDKRSPGWGKRDSTDFLDNSLDKRRPGWGKRSDILDRIAMLKRRPGWGKRTVSDFSDLDKRRPGWGKRDSEGIDMEIRAPGWGKRNSIDDREFSLYDKRRPGWGKRSDNIELETRRPGWGKRAPGWGKRSMDVCLSMKEKAEYLINSAIEIETEYASLCGIRDSRS

**Gene structure analyses:**

**RGWamide/APGWamide**

**Mizuhopecten yessoensis (Mollusca)**

>M. yessoensis APGWamide AXN93473.1 (precursor)

MDSLTVGILSFILTFNVLSASSDTDIMDKRRPGWGKRDGQSEESTIEEFGEPDKRSPGWGKRDSTDFLDNSLDKRRPGWGKRSDILDRIAMLKRRPGWGKRTVSDFSDLDKRRPGWGKRDSEGIDMEIRAPGWGKRNSIDDREFSLYDKRRPGWGKRSDNIELETRRPGWGKRAPGWGKRSMDVCLSMKEKAEYLINSAIEIETEYASLCGIRDSRS

>M. yessoensis APGWamide AXN93473.1 (splice pattern)

MDSLTVGILSFILTFNVLSASSDTDIMDKRRPGWGKRDGQSEESTIEEFGEPDKRSPGWGKRDSTDFLDNSLDKRRPGWGKRSDILDRIAMLKRRPGWGKRTVSDFSDLDKRRPGWGKRDSEGIDMEIRAPGWGKRNSIDDREFSLYDKRRPGWGKRSDNIELETRRPGWGKRAPGWGKRSMDVCLSMKEKAEYLINSAIE—(phase 0)---28339 ------IETEYASLCGIRDSRS

>M. yessoensis APGWamide MH045205.1 (mRNA)

ATGGATTCTTTAACCGTAGGAATTCTAAGCTTTATTTTAACATTCAACGTGCTATCTGCATCCAGTGATACAGACATTATGGATAAGCGCAGACCCGGATGGGGAAAACGAGATGGTCAGTCGGAGGAGTCCACGATTGAAGAGTTCGGAGAACCCGACAAACGTAGTCCTGGATGGGGAAAACGGGATTCCACAGATTTCTTAGATAATTCTCTTGATAAACGTCGCCCTGGATGGGGCAAACGTAGTGACATTTTGGACAGGATCGCGATGCTGAAAAGGAGACCCGGTTGGGGAAAAAGAACCGTATCGGACTTTTCCGATTTAGATAAAAGAAGGCCAGGATGGGGAAAAAGGGATTCCGAAGGAATCGACATGGAAATAAGGGCACCAGGATGGGGGAAAAGGAACTCCATTGATGACAGAGAATTTTCTCTATATGACAAAAGGCGACCAGGATGGGGGAAACGTTCCGACAATATAGAGTTAGAAACACGAAGACCCGGATGGGGAAAACGAGCACCCGGATGGGGAAAAAGATCCATGGATGTTTGTCTGTCTATGAAGGAAAAAGCTGAATATTTAATAAACTCCGCTATTGAGATTGAGACAGAATACGCTTCACTCTGTGGAATTCGTGATTCAAGGTCATGA

Genome: NEDP02005386.1

**Lingula anatina (Brachiopoda)**

>L. anatina GWamide XP_013419038.1 (precursor)

MKSFLVIVELLVYSLLFSSCYVQGGDRYRRRMGWGKRSEPQVSDISYLLSAVDSLSQNPPGNNGDDMDRPNRGWGKRFSGSDMASTMDKRGWGKRSTTLDLEEVKRAMGWGKRSFFNESPVRRAMGWGKRERAPAGWGKRNQILVPELKKGLGWGKRILKSDDVADVEDAPQMDYNRRFLPGNSSASATDLSSPVSNKESTDIRDKSAQCQNNIEAFSRMLDMFVKILTYESCDLEREVISKMLAQVRMSLLID

>L. anatina GWamide XP_013419038.1 (splice pattern)

MKSFLVIVELLVYSLLFSSCYVQGGDRYRRRMGWGKRSEPQVSDISYLLSAVDSLSQNPPGNNGDDMDRPNRGWGKRFSGSDMASTMDKRGWGKRSTTLDLEEVKRAMGWGKRSFFNESPVRRAMGWGKRERAPAGWGKRNQILVPELKKGLGWGKRILKSDDVADVEDAPQMDYNRRFLPGNSSASATDLSSPVSNKESTDIRDKSAQCQNNIEAFSRMLDMFVK-(phase 0)----2644----ILTYESCDLEREVISKMLAQVRMSLLID

>L. anatina GWamide XM_013563584.1 LOC106179816 (mRNA)

CCAGTGAGGAAAAATGAAGTCGTTTTTGGTAATAGTCGAACTGCTTGTATACTCACTTCTCTTCTCATCTTGCTATGTGCAAGGCGGCGACAGATATAGAAGAAGAATGGGATGGGGAAAACGCAGCGAGCCACAAGTGAGCGATATTTCGTACCTGCTTTCGGCTGTCGATTCACTGTCTCAAAATCCTCCTGGCAATAACGGCGATGATATGGATCGGCCAAATCGTGGCTGGGGAAAACGCTTCTCCGGCAGTGACATGGCAAGCACGATGGACAAACGAGGGTGGGGCAAAAGATCCACTACTCTCGACCTGGAAGAGGTGAAACGAGCGATGGGATGGGGTAAACGCTCTTTTTTCAATGAGTCCCCGGTCCGACGCGCTATGGGGTGGGGGAAGCGAGAAAGAGCACCAGCAGGGTGGGGAAAACGTAATCAAATTTTGGTCCCGGAGTTAAAAAAAGGCTTAGGCTGGGGAAAACGTATTCTCAAAAGTGACGATGTCGCAGATGTAGAGGACGCCCCTCAGATGGACTACAACAGGCGTTTTCTCCCAGGAAATTCCTCCGCATCAGCAACAGACTTATCTTCTCCAGTTTCAAATAAAGAGTCCACGGACATAAGGGACAAATCAGCACAGTGTCAAAACAATATAGAAGCATTCTCTCGGATGCTGGACATGTTTGTGAAGATATTGACCTATGAATCGTGTGATTTAGAGAGGGAAGTTATTAGTAAAATGCTTGCCCAAGTACGAATGAGCTTGTTAATAGACTAAAAACAGGTTCCCAAGGATATGTCGTATTTCCCATCACCTTCAGCCTTGTGGTAAATTATTTTTAATCTTAAAATCTACGAAACGAATGGTCTCAAACAGTGGTGTACTTGAAACACTATTACCTCGCTCTCAAGAGTATTTTTCAAGACTTCTTATGTGCATGACTGTACACTCGCTGTAATTAGCACAG

Genome: NW_019775415.1

**Notospermus geniculatus (Nemertea)**

>N. geniculatus GWamide g26977.t1 (precursor)

MKFYIAITLVCLLVFDIFASAAAAENADQDLDKRAPGWGKRGWGKRDDESDASLDDYLKRGWGKRGWGKRGWGKRGWGKRDDTPSCDDLNQGVLLFIYKAIEAEAQRITACSSQEN

>N. geniculatus GWamide g26977.t1 (splice pattern)

MKFYIAITLVCLLVFDIFASAAAAENADQDLDKRAPGWGKRGWGKRDDESDASLDDYLKRGWGKRGWGKRGWGKRGWGKRDDTPSCDDLNQGVLLFIYKAIE-(phase 0)--------12912 -------AEAQRITACSSQEN

>N. geniculatus GWamide scaffold1472.g26977.t1 (mRNA)

Atgaagttctacatcgccattacccttgtctgcctcctagtattcgatatcttcgccagtgccgcagcggccgaaaatgccgaccaggaccttgacaaacgtgccccgggatggggcaagaggggctggggcaagcgggacgatgaatctgacgcctcattggatgactacctgaagcgaggctggggcaagagaggctggggcaaacgtggctggggaaagcgaggatggggcaagagggacgacactccatcatgtgacgatttaaaccagggtgtactcctctttatttacaaagctattgaggccgaggctcaaagaatcaccgcatgttcaagccaagaaaactaa

Genome: NMRB01000859.1

**Capitella teleta (Annelida)**

>C. teleta RGWamide EY540104.1 CAPTEDRAFT_100873 (precursor)

MLLNHLILLSLALAFIPAVSFASDELVDEEASEMDKRRGWGKRSDSEMDKRRGWGKRSFDEDSEEVAKRRGWGKRSEAINLDEVEESIAEMDKRRGWGKRSELEAMQSEFAKRRGWGKRADDMEILKRRGWGKRSEDANEMDKRRGWGKRSLEAEKRRGWGKRSPKIEEFSEIVDCQEVKEKIFYHVSLALQLDAQRANICPAEK

>C. teleta RGWamide EY540104.1 CAPTEDRAFT_100873 (splice pattern)

MLLNHLILLSLALAFIPAVSFASDELVDEEASEMDKRRGWGKRSDSEMDKRRGWGKRSFDEDSEEVAKRRGWGKRSEAINLDEVEESIAEMDKRRGWGKRSELEAMQSEFAKRRGWGKRADDMEILKRRGWGKRSEDANEMDKRRGWGKRSLEAEKRRGWGKRSPKIEEFSEIVDCQEVKEKIFYHVSLALQ-(phase 0)--7181-----LDAQRANICPAEK

>C. teleta RGWamide EY540104.1 (mRNA)

CGGTCCGGATTCCCGGGATTGAGATCGACACTCAGCGACCGTCCAAAAAGTTTCTGGAGTGTTATATCTCTGGCAATGCTGCTGAACCATTTGATTCTACTGAGCTTGGCGCTCGCCTTCATCCCGGCCGTCTCGTTTGCCAGCGATGAACTAGTGGACGAGGAAGCGTCCGAGATGGACAAGCGACGAGGGTGGGGAAAACGCAGCGACAGTGAAATGGACAAACGAAGAGGTTGGGGCAAGAGAAGTTTCGACGAGGATTCCGAAGAGGTCGCAAAGAGGAGAGGTTGGGGGAAGAGGTCCGAAGCCATTAATTTGGACGAGGTGGAGGAGTCAATAGCTGAAATGGACAAGCGAAGAGGCTGGGGGAAGCGGTCCGAATTGGAAGCCATGCAGTCTGAATTCGCTAAACGACGAGGCTGGGGCAAGAGAGCCGATGACATGGAAATTTTGAAGAGGAGAGGATGGGGTAAGCGAAGTGAAGATGCCAATGAAATGGACAAGAGACGAGGCTGGGGAAAGAGATCATTGGAAGCGGAGAAGAGGAGAGGTTGGGGCAAACGCAGTCCTAAAATCGAGGAATTCAGCGAAATCGTCGATTGTCAGGAAGTCAAGGAGAAGATCTTTTACCACGTCAGTCTTGCTCTCCAGCTGGATGCACAAAGAGCTAACATTTGCCCCGCCGAGAAATAAAGTCGACATGTGCCTCGTTTGAACGTGGTGCAGTGCTGCCACAAAAACATTAAAAAACAAAAAACAAAAGAGATTCGTTCTTACGCATGTGACGTCACAACTTCTTCTCGTCATCTTCTATTGCTCTGCCAAGAATCAAGGCTCGCAATCTGATTGGTAGATGGCACTTCCGGTCCATGAACTTCCTGTAATCAAAATTCAAATCTATTAAAATTGTATGCGTGTGA

Genome: AMQN01004918.1

**Pleurin**

**Mizuhopecten yessoensis (Mollusca)**

>M. yessoensis pleurin AXN93514.1 (precursor)

MSPHSILTLLTTFAIIQLAKAIFYTNKEGNDFPRIGKRRHLYQSEAWSRDSVPSDGLPGETDRQMEPMYNKDEGEIPITSRREHQNPMSSYSKAMYYRALGLNGYRAFSKNKVQASPNNGIRGTRTYY

>M. yessoensis pleurin AXN93514.1 (splice pattern)

MSPHSILTLLTTFAIIQLAKAIFYTNKEGNDFPRIGKRRHLYQSEAWSRDSVPSDGLPGETDRQMEPMYNKDEGEIPITSRREHQNPMSSYSKAMYYRALGLNG-(phase 1)-320bp----YRAFSKNKVQASPNNGIRGTRTYY

>M. yessoensis pleurin MH045246.1 (mRNA)

ATGTCACCGCATTCAATCCTTACTCTCTTAACAACATTCGCCATCATACAGCTAGCCAAGGCCATATTCTACACTAACAAGGAGGGCAATGATTTTCCTAGAATTGGTAAACGGCGACATCTGTATCAATCTGAAGCTTGGTCACGTGACTCGGTCCCCTCTGACGGTCTTCCTGGAGAGACAGACCGACAGATGGAGCCTATGTATAATAAAGACGAAGGGGAAATACCAATCACTTCCAGACGTGAACACCAAAATCCAATGTCCAGTTACTCTAAAGCTATGTACTATCGGGCACTCGGCCTAAACGGTTACCGTGCATTTTCCAAAAACAAGGTCCAGGCGTCTCCAAACAATGGAATACGAGGAACACGTACATATTACTAA

Genome: NW_018407522.1

**Lingula anatina (Brachiopoda)**

>L. anatina pleurin comp14311_c0_seq1 (precursor)

MILKFTCYVGLACVLLSALQGTYCFYFTDSKEHDYPRIGRRNQIVSTKAHFDFENKGLVNSNIQLPGSSDVIFANRNIEALLQRLSVPMVSYSDASRLSDAWLRRRRSLLSYQKSQ*

>L. anatina pleurin comp14311_c0_seq1 (splice pattern)

MILKFTCYVGLACVLLSALQGTYCFYFTDSKEHDYPRIGRRNQIVSTKAHFDFENKGLVNSNIQLPGSSDVIFANRNIEALLQRLSVPMV-(phase 1)-2100bp-----SYSDASRLSDAWLRRRRSLLSYQKSQ

>L. anatina pleurin comp14311_c0_seq1 (mRNA)

ATGATTCTCAAATTCACCTGTTACGTTGGACTTGCGTGCGTCCTTTTGTCTGCACTCCAAGGGACCTACTGCTTTTACTTCACAGATAGCAAAGAACATGATTATCCGCGCATAGGAAGACGGAACCAAATTGTTTCAACGAAAGCTCATTTTGATTTCGAAAACAAAGGGCTGGTAAATTCTAATATACAGCTTCCCGGCAGCTCCGATGTTATATTTGCCAACAGAAACATAGAGGCACTGCTTCAAAGACTGAGCGTTCCAATGGTATCCTATAGTGACGCCAGCAGATTAAGCGATGCGTGGCTACGTCGGAGGAGAAGTCTGTTGTCATACCAGAAGTCACAGTAA

Genome: NW_019775924.1

**Notospermus geniculatus (Nemertea)**

>N. geniculatus pleurin g3565.t1 (precursor)

MESRLVLAQALLVVCLVHFAHGIFFTSSKENEYPRIGRRNFVARADSTDLNDSRDLYEQRLPVLAKALYRELSARGVKPYQFKGMRTENGADDFDTKTRHILGED

>N. geniculatus pleurin g3565.t1 (splice pattern)

MESRLVLAQALLVVCLVHFAHGIFFTSSKENEYPRIGRRNFVARADSTDLNDSRDLYEQRLPVLAKALYRELSARG-(phase 1)-3981bp----VKPYQFKGMRTENGADDFDTKTRHILGED

>N. geniculatus pleurin scaffold63.g3565.t1 (mRNA)

Atggagtcacgcctggttttagcacaagccctcctggtcgtctgcttggtgcatttcgcacatggaatcttcttcacgagttccaaggaaaatgaatatcctcggattggcagacgtaatttcgtagctcgtgcggactctactgatttaaacgattcgagagatttatacgaacaaagattgccagttttggcgaaagctttatacagggaactttcagctcggggtgtaaaaccttaccaatttaagggaatgagaaccgaaaacggagctgacgatttcgacacgaagacgaggcatatcctcggcgaggactaa

Genome: NMRB01000063.1

**CCWamide/Agatoxin**

**Lottia gigantea (Mollusca)**

>L. gigantea CCWa XP_009055999 LOTGIDRAFT_161866 (precursor)

MNVDHSVVRYTDTRFCGNECRSWCNTESPFHCHRYSMRNYNDFDALDKLKTIKRRVSFRRNQQVEPRTCRGWRAPCIPWTSEPGEACCTSANLVCRCNLWMQNCRCVGRTWG

>L. gigantea CCWa XP_009055999 LOTGIDRAFT_161866 (splice pattern)

MNVDHSVVRYTDTRFCGNECRSWCNTESPFHCHR-(phase 1)-3591-----YSMRNYNDFDALDKLKTIKR-(phase 2)—927----RVSFRRNQQVEPRTCRGWRAPCIPWTSEPGEACCTSANLVCRCNLWMQNCRCVGRTWG

>L. gigantea CCWa XM_009057751.1 (mRNA)

ATGAATGTAGATCATAGTGTAGTACGATACACCGATACAAGGTTCTGTGGCAATGAATGTAGATCATGGTGTAATACTGAATCCCCTTTTCACTGCCATCGTTATTCTATGAGAAATTATAATGATTTTGATGCCTTGGATAAACTAAAAACAATTAAAAGACGAGTTTCATTTAGGAGAAATCAACAAGTAGAACCAAGAACATGTAGAGGATGGAGGGCACCGTGTATACCTTGGACATCTGAACCAGGAGAAGCATGTTGTACTAGTGCAAATTTAGTATGTCGATGTAATTTATGGATGCAGAATTGTCGATGTGTCGGTCGAACCTGGGGATGA

Genome: AMQO01002781.1

**Crassostrea virginica (Mollusca)**

>C. virginica CCWa-1 XP_022294698.1 CCW1 amidated (precursor)

MTFFVAGFWIFLVSLNIGTCMYYADYDDDYYNQNKALDLFKSMDRRNYYLSQPTCNGWNRHCLPWSSQIQHSCCGGLSCKCNLWGQNCRCTTKLWGR

>C. virginica XP_022294698.1 LOC111104838, isoform X2 (splice pattern)

MTFFVAGFWIFLVSLNIGTCMYYAD-(phase 1)--4934—YDDDYYNQNKALDLFKSMDR-(phase 2)--4151-----RNYYLSQPTCNGWNRHCLPWSSQIQHSCCGGLSCKCNLWGQNCRCTTKLWGR

>C. virginica XM_022438990.1 LOC111104838, transcript variant X2 (mRNA)

TCTGTAAGGATGACTTTTTTCGTAGCTGGATTTTGGATTTTCCTTGTTTCTCTCAATATTGGAACTTGCATGTATTATGCAGATTACGATGATGATTACTACAACCAGAACAAAGCCCTAGATTTATTCAAGTCAATGGATAGACGTAACTATTACCTGAGTCAGCCCACGTGCAACGGTTGGAACCGCCATTGCCTCCCTTGGTCATCGCAGATTCAGCACAGTTGCTGTGGGGGGCTTTCATGCAAGTGTAACCTCTGGGGTCAGAATTGTAGGTGTACGACAAAGTTATGGGGGCGCTGATTGGACAAGGCGCGTCACGTGACCGAAGAGACACGATTTCTAGTGTCATGGATTTCATTTCAGAACTACCCATTTTATCGTAAAAATAAAGTAAACATTGTATTACACACAC

>C virginica XM_022438989.1 LOC111104838, transcript variant X1 (mRNA)

GTTCTGTAAGGATGACTTTTTTCGTAGCTGGATTTTGGATTTTCCTTGTTTCTCTCAATATTGGAACTTGCATGTATTATGCAGATTACGATGATGATTACTACAACCAGAACAAAGCCCTAGATTTATTCAAGTCAATGGATAGACGTAACTATTACCTGAGTCAGCCCACGTGCAACGGTTGGAACCGCCATTGCCTCCCTTGGTCATCGCAGATTCAGCACAGTTGCTGTGGGGGGCTTTCATGCAAGTGTAACCTCTGGGGTCAGAATTGTAGGTGTACGACAAAGTTATGGGGGCGCTGATTGGACAAGGCGCGTCACGTGACCGAAGAGACACGATTTCTAGTGTCATGGATTTCATTTCAGAACTACCCATTTTATCGTAAAAATAAAGTAAACATTGTATTACACACAC

Genome: MWPT03000008.1

**Notospermus geniculatus (Nemertea)**

>N. geniculatus CCWa1 g41399.t1 (precursor)

MLPNSVVDDSGGTCSQTKKLFEKCTQGEYGDDQLEVFRDAAKRAFYNFKCATSTCRPRSMEPTLRCCPGMRCKCGILWNAGQCKCTNYGYGR

>N. geniculatus CCWa1 g41399.t1 (splice pattern)

MLPNSVVDDSGGTCSQTKKLFEKCTQ—(phase 1)--------1188-----GEYGDDQLEVFRDAAK-(phase 2)------4513-----RAFYNFKCATSTCRPRSMEPTLRCCPGMRCKCGILWNAGQCKCTNYGYGR

>N. geniculatus CCWa-1 scaffold5958.g41399.t1 (mRNA)

atgctcccgaacagtgtagtggatgacagcggcggcacctgctcacaaactaagaaactatttgaaaaatgtacacagggtgaatacggtgatgaccaactagaagtcttcagagatgccgccaagagagccttttacaatttcaaatgtgcgacctcgacctgccgaccgcgatctatggaaccaaccctccgatgctgcccaggaatgcgatgcaagtgcggcattctgtggaacgccggccagtgcaagtgcaccaactacggctacggacgatga

>N. geniculatus CCWa-2 scaffold52.g3053.t1.p1 (precursor)

MKLYIVACLFVVCALHFVASYPYGEEKAKKSADNQEGEYGDDQLEVFRDAANDLETPPERRAFYNFKCATSTCRPRSMEPTLRCCPGMRCKCGILWNAGQCKCTNYGYGR

>N. geniculatus CCWa-2 scaffold52.g3053.t1.p1 (splice pattern)

MKLYIVACLFVVCALHFVASYPYGEE-(phase 1)--65988---KAKKSADNQE-(phase 1)----20250--GEYGDDQLEVFRDAAN--(phase 2)-1532---DLETPPER(phase 2)--2981-RAFYNFKCATSTCRPRSMEPTLRCCPGMRCKCGILWNAGQCKCTNYGYGR

>N. geniculatus CCWa-2 scaffold52.g3053.t1 (mRNA)

atgaagctatatattgtggcttgcctattcgtagtctgcgcactacacttcgttgccagttacccatatggagaagagaaggctaaaaagagtgctgacaatcaagagggtgaatacggtgatgaccaactagaagtcttcagagatgccgccaatgaccttgagactcccccagaaaggagagccttttacaatttcaaatgtgcgacctcgacctgccgaccgcgatctatggaaccaaccctccgatgctgcccaggaatgcgatgcaagtgcggcattctgtggaacgccggtcaatgcaagtgcaccaactacggctacggacgatga

(original transcript has an incorrect N-terminus – the correct gene starts with the second Methionine)

Genome: NMRB01000052.1 (*The two CCWamides 1 & 2 are derived from the same gene with alternative splicing which changes the amino acids in the edges of one of the exon transcripts.)*

>N. geniculatus CCWa-like 3 scaffold6.g467.t1.p1 (precursor)

MDSRWIYTVLIILLLVNVSVCFVRREDDKRSEEKESATCRKKKEFCELDNPGLTCCTGLMCSCGLYDYCRCSDPMEWNEN

>N. geniculatus CCWa-like 3 scaffold6.g467.t1.p1 (splice pattern)

MDSRWIYTVLIILLLVNVSVCFVRRE-(phase 1)-314—DDKRSEEKES-(phase 2)-1278—ATCRKKKEFCELDNPGLTCCTGLMCSCGLYDYCRCSDPMEWNEN

>N. geniculatus CCWa-like 3 scaffold6.g467.t1 (mRNA)

Atggattcgaggtggatttatacagttctcatcatcctgttgctagtcaacgtgtctgtttgttttgtgagacgggaagacgacaagcgatctgaagaaaaagaaagcgccacctgccggaaaaagaaagaattctgcgaacttgacaacccaggtcttacctgttgcaccgggctgatgtgctcgtgtggtttatatgactactgccgatgctcggatccgatggaatggaacgaaaattga

Genome: NMRB01000006.1

**Phoronis australis (Phoronis)**

>P. australis CCWa TRINITY_DN320441_c3_g7_i1 (precursor)

METGTSRVVLALVLSLHIALSWSHLVFDSKENELALNKLRAAKRVVWRYRVKSCASWNDYCDPWPDESSQDRLTRSFTCCDNMICKCNLWAQNCRCKSRIWGR

>P. australis CCWa TRINITY_DN320441_c3_g7_i1 (splice pattern)

METGTSRVVLALVLSLHIALSWSHLVFD-(phase 1)---7560-----SKENELALNKLRAAKR-(phase 2)---2201----VVWRYRVKSCASWNDYCDPWPDESSQDRLTRSFTCCDNMICKCNLWAQNCRCKSRIWGR

>P. australis_TRINITY_DN320441_c3_g7_i1 (mRNA)
GCTTCACTCAATTAACGGCTCTAGATTAGTTATGCGAGTGTGCTGCTGGCATTGTTAAGTTTACTGAGGGTTTCCACTATCGGGGGACACAGTTTTGCAGACGTCTATTGTTTTATGCCATTGGAAGCGCAGTGGACCCAAGAAACATGGAAACTGGAACAAGTCGGGTAGTTTTAGCTCTGGTGTTAAGCCTGCACATTGCACTCAGTTGGTCACATCTAGTCTTTGATTCAAAAGAAAATGAATTAGCTTTAAACAAGTTGAGGGCGGCAAAGCGGGTGGTATGGAGATACCGTGTGAAGTCGTGTGCGTCTTGGAACGACTATTGCGACCCCTGGCCGGATGAAAGCAGCCAGGATAGGCTAACGAGGTCGTTCACGTGTTGCGATAACATGATATGCAAATGTAATTTGTGGGCGCAAAACTGTAGGTGTAAGTCCCGGATATGGGGACGCTGATAACATGCGTCTAAGGATAAGTTTACACGTGCCATTGGGGAGCTATATTAGAAATAGAGCAATCTCCATATGAGAGGGACAGTTTCCGTGGTACGTTTTCACCAGAGGCGTCAACATTGTAAAAATTGCGCAAAAGAATGAATTTGTTATTTTAACACTTGTACGCTACATCTTTACTGCGATGCCGTTTTAATTAGCTGTCGATGAAACATCAATTGTGTTCCTGGAAGCAATAACTTTGGTAACTTCAGTTCCAAGTGGCCACGCCATGAAAAAGACTGAAGAACGATTGAGTGCCTGACATCTGTACAGTATGTGCTTCAGTCGAACACTTGTTGTTTAACAAACACGAGCAATTCCGAGAGTTGCCAGATTTCTCACAACTCTTCTTCTAGACTTCTTT

Genome: NMRA01000043.1

**Tribolium castaneum (Hexapoda)**

>T. castaneum ALP EFA01531.2 TcasGA2_TC007091 (precursor)

MKYTWLVLAACMVLVLAELLPGAAAGPYLDDDEGLPSDDDYTENAIDRLLQSAQKRSSLIYLFRRACVRRGGNCDHRPNDCCYNSSCRCNLWGSNCRCQRMGLFQKWG

>T. castaneum ALP EFA01531.2 TcasGA2_TC007091 (splice pattern)

MKYTWLVLAACMVLVLAELLPGAAAGPYLDDD-(phase 1)-5473--—EGLPSDDDYTENAIDRLLQSAQK—(phase 2)---1004----RSSLIYLFRRACVRRGGNCDHRPNDCCYNSSCRCNLWGSNCRCQRMGLFQKWG

>T. castaneum ALP XM_008194546.2 LOC656848, transcript variant X1 (mRNA)

TTCGAATTATTTCACGTATCATGAAGTACACTTGGTTGGTATTAGCGGCCTGCATGGTGTTGGTTCTCGCGGAGTTGCTCCCGGGGGCCGCTGCAGGACCCTATCTAGATGACGACGAAGGCCTCCCTTCGGACGACGACTACACGGAAAACGCAATCGATCGTTTGTTACAATCTGCTCAGAAACGTTCTTCCTTAATTTACCTTTTCAGACGAGCCTGTGTGAGAAGAGGAGGGAACTGCGACCATCGACCGAACGACTGCTGCTATAACAGTTCGTGTCGTTGCAATTTGTGGGGCTCTAATTGCAGATGTCAACGAATGGGACTGTTCCAGAAATGGGGCTAGACCTTGAACTAATTTAAGCTGAAACGGAGGGCCTCGAGCGGCGGTCGCCGGATGAATTATTTTCTTTACGAAAAGAACTGATTTTTGTATTAATTGCGGCTCCGTTTGAGGCTTTCTCCAAAAACCGACAATTGGCGCGATTACGGATACGATTAATGCATGGTTTAAGTTCCTTTGGTTTTGTTTCGTCGAACCCAAACATTGCTTTGTGAAGTCGATAATAATTAAAACAATAAAAAAATGAACGAATTACAATTTATTTTTGATACTTTGCTTATACAGCGTTCTAGTTTTGAGGCTTCTTTCATTTTAGGTTTCACTGATGTTGTCATCGACATCTTTTGACAACGATACTTTTGACTAAAAGAAAGAATTTCTGTGTGCGAAAATTCCAATTCGGCACAAATTATCGAATCGTGACGATGAGTGCGACATCTGACGGTGATCTGATTCGAAGTTCGTAATTTTTGCCGAATTACACAATCAGCTGCTCTCCCATTAAAGGTTTTTCCAGCCACACCCTTAATGGGAATGCATGTCTGACGTTATCTTAGGTAATTTCTATAATATGACACTATATTGTAATTATGTATGTATGTATAGATGTAATGTCATTAAGGCGACATCTGTGGTGCTAGTGATAAGGCCGCAGATTCACCATCTAACATTGTTGTTAGTGATTTGAATCCTATATAAATATGTTTCTATGTGAAAGGGTACAACATTCAAATGATTCTGTCAATACTAAATATATCAATTGGCCTA

Genome: AAJJ02001753.1

**Centruroides sculpturatus (Arachnida)**

>C. sculpturatus U8-agatoxin-Ao1a-like-1 XP_023230485.1 (precursor)

MNLTLLMLVFCLLICSSLSTPYYGSDNSLDDYNDGLTRYLLYTRKRSCIRRGGFCDNRPNDCCFNSSCRC

NLWGTNCRCQRPGLFQKWGK

>C. sculpturatus U8-agatoxin-Ao1a-like-1 XP_023230485.1 (splice pattern)

MNLTLLMLVFCLLICSSLSTPYYGSD-(phase 1)-149—NSLDDYNDGLTRYLLYTRK-(phase 2)---626-RSCIRRGGFCDNRPNDCCFNSSCRCNLWGTNCRCQRPGLFQKWGK

>C. sculpturatus U8-agatoxin-Ao1a-like-1 XM_023374717.1 LOC111630597 (mRNA)

AAAACATGAACTTGACTTTGTTAATGTTGGTCTTCTGCCTGTTGATTTGTTCGTCTTTAAGCACTCCATATTATGGATCTGATAACTCTCTAGACGACTACAACGATGGACTGACTCGATACTTACTCTACACTAGAAAGAGAAGTTGCATCAGACGCGGTGGTTTCTGCGACAATAGGCCAAATGATTGTTGTTTCAACTCTTCATGTCGCTGCAACCTCTGGGGAACCAACTGTCGATGTCAGAGACCGGGACTCTTTCAGAAATGGGGAAAGTAAAATGCCGTTTGCAGGTTGTTTTTGAAGAAGAGGAAGAAGAAAAGCCGAACTTGGTGTAGACGGAGCCCTGTTGTTTTGGGCATCGACACTACATTGTGGACATTATAGATAATATATGTTTGATAAATATATATTTCCAACTCCACAATGTAACCTTCCAAAGGCGGAATTCTTTTGGAAACACGTGGTGTTAATCCATTCATTTTAACCCGTTGTGCTATGTAATTTGTTGTGTTATATTCCCTTTAAAGTATATAAAGGCCGTAAGCAGAATAAAAAAGTAACTTTGTGCA

Genome: NW_019384857.1

>C. sculpturatus U8-agatoxin-Ao1a-like-2 XP_023230486.1 (precursor)

MKHSAVISIVLILILAQSLNVMSYYGNEASLLDDYNDGLARFLFFTRKRSCIRRGGSCDHRPHDCCFSSSCRCNLWGTNCRCQRPGLFQKLGK

>C. sculpturatus U8-agatoxin-Ao1a-like-2 XP_023230486.1 (splice pattern)

MKHSAVISIVLILILAQSLNVMSYYGNE-(phase 1)--402---ASLLDDYNDGLARFLFFTRK-(phase 2)--1804-RSCIRRGGSCDHRPHDCCFSSSCRCNLWGTNCRCQRPGLFQKLGK

>C. sculpturatus U8-agatoxin-Ao1a-like-2 XM_023374718.1 LOC111630598 (mRNA)

GACGAGTATGAAACATTCTGCCGTTATTAGTATTGTTCTGATACTTATTTTGGCTCAGAGTTTAAATGTTATGTCATATTATGGTAACGAAGCCAGTTTATTGGATGATTACAACGATGGACTCGCTCGTTTTCTATTCTTTACGAGAAAGAGAAGCTGCATAAGACGTGGTGGAAGTTGCGATCATCGACCCCATGATTGCTGTTTTAGTTCTTCGTGTCGATGTAATCTTTGGGGTACCAACTGTAGATGCCAAAGACCAGGATTGTTCCAGAAATTGGGAAAATAAATTTTCTTTGGATAATTATGTTTATCTTACCAAACAACTTCTGCTTATTGTATAATTTATTTTCCAAATAAATTTATTCTGTGTGCTCAAGGCACTTATTACAAAGACAGTAATATATATAAATATATACTTCTGAGCTTCCCAAGAAAGAAATTTTAACGTTTTTGAAAAATATATTAATAAATATACAAAAAAAAATCTTTTCGAACTTAAGAAATATTTATTGTTAATACGTTTTTGAAAAATCATAATATTCGTTATAATGTTAATCAAGTTATTTAAG

Genome: NW_019384857.1

>C. sculpturatus U8-agatoxin-Ao1a-like-3 XP_023230478.1 (precursor)

MNCVSILVIFGLVLLADAILASPYTPTDDEAVEDYSDRLENLILNAEKRNCIRRGKSCDNKPNGCCENSSCRCNLWGTNCRCQRAGLFQRWGK

>C. sculpturatus U8-agatoxin-Ao1a-like-3 XP_023230478.1 (splice pattern)

MNCVSILVIFGLVLLADAILASPYTPTD-(phase 1)--86-DEAVEDYSDRLENLILNAEK-(phase 2)---184---RNCIRRGKSCDNKPNGCCENSSCRCNLWGTNCRCQRAGLFQRWGK

>C. sculpturatus U8-agatoxin-Ao1a-like-3 XM_023374710.1 LOC111630593, transcript variant X1 (mRNA)

TCAACTAAAGATATGAATTGCGTTTCGATCCTGGTCATTTTTGGATTGGTTCTCTTGGCTGATGCCATTCTAGCAAGTCCATATACACCGACCGATGATGAAGCGGTAGAAGATTACAGCGATCGATTGGAAAATTTGATTTTGAATGCAGAAAAACGAAATTGTATTCGAAGAGGCAAATCGTGCGATAACAAACCCAACGGATGCTGCGAAAACTCATCGTGTCGTTGCAATTTGTGGGGTACCAATTGCCGTTGCCAAAGAGCAGGCCTCTTCCAACGATGGGGTAAATAAGCCAGCGTACTTTTGTGTCTGTCAACCAGTATACAAGATACATGTCTATCCATAAAGCATTTTAGAAGATCGTTTAATATCGGATATATATATAATATATATATATACATATGTATATGTAATTATTGTTAAAAATTGTTCCGAAAATGTTACTCATGTTTAAAAACGTTAAAAAATCAAAAACATTTACTTTTCCTTTAAAATATTTTCCTAAACAATATTGTACAACGATGTAGAATTAGATATTCCACATGTTAATAAACAAAATGTAATAAAATAGTATTATTTTTTCCTCATATTTTAAGTAGTAAAGCAATTATTTTCTGTAAAATCTCTATAATTTTAATATAAAAATAATAGAAGATATAATAAATCAATTTTGTTCAATTAATTGTTTAAATTAAACAATAAAGTTCACATTTAAAATGG

Genome: NW_019384857.1
