## Supplementary figures and images for "Nemertean, brachiopod and phoronid neuropeptidomics reveals ancestral spiralian signalling systems"

### Supplementary_Material_10_Vasotocin_trees.pdf

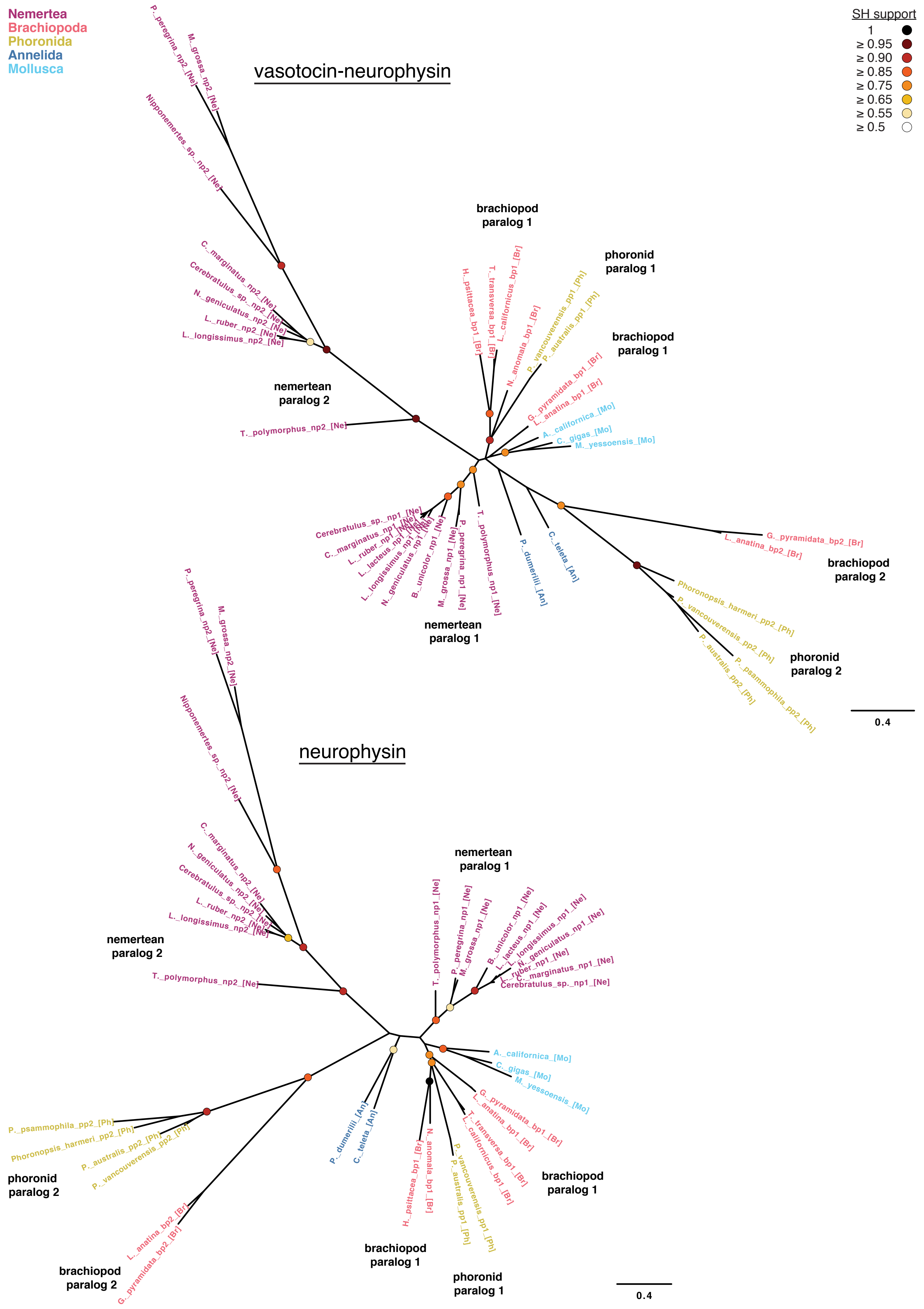
